# A pair of isoleucyl-tRNA synthetases in Bacilli fulfill complementary roles to keep fast translation and provide antibiotic resistance

**DOI:** 10.1101/2022.02.09.479832

**Authors:** Vladimir Zanki, Bartol Bozic, Marko Mocibob, Nenad Ban, Ita Gruic-Sovulj

## Abstract

Isoleucyl-tRNA synthetase (IleRS) is an essential enzyme that covalently couples isoleucine to the corresponding tRNA. Bacterial IleRSs group in two clades, *ileS1* and *ileS2,* the latter bringing resistance to the natural antibiotic mupirocin. Generally, bacteria rely on either *ileS1* or *ileS2* as a standalone housekeeping gene. However, we have found an exception by noticing that *Bacillus* species with genomic *ileS2* consistently also keep *ileS1,* which appears mandatory in the family *Bacillaceae*. Taking *Priestia (Bacillus) megaterium* as a model organism, we showed that PmIleRS1 is constitutively expressed, while PmIleRS2 is stress-induced. Both enzymes share the same level of the aminoacylation accuracy. Yet, PmIleRS1 exhibited a twofold faster aminoacylation turnover (*k*_cat_) than PmIleRS2 and permitted a notably faster cell-free translation. At the same time, PmIleRS2 displayed a 10^4^-fold increase in its *K*_i_ for mupirocin, arguing that the aminoacylation turnover in IleRS2 could have been traded-off for antibiotic resistance. As expected, a *P. megaterium* strain deleted for *ileS2* was mupirocin-sensitive. Interestingly, an attempt to construct a mupirocin-resistant strain lacking *ileS1*, a solution not found among species of the family *Bacillaceae* in nature, led to a viable but compromised strain. Our data suggests that PmIleRS1 is kept to promote fast translation, whereas PmIleRS2 is maintained to provide antibiotic resistance when needed. This is consistent with an emerging picture in which fast-growing organisms predominantly use IleRS1 for competitive survival.

## Introduction

Bacteria respond to various stress conditions by changes in transcription and/or translation [1, 2]. Important players in the expression of genetic information are aminoacyl-tRNA synthetases (aaRSs) [3, 4]. These housekeeping enzymes supply ribosomes with aminoacylated tRNAs (aa-tRNA) and secure the optimal speed of translation and cellular fitness under normal and starvation conditions [5]. Aminoacylation is a two-step reaction localized within the aaRS synthetic site [3]. The amino acid is firstly activated at the expense of ATP to form an aminoacyl-adenylate intermediate (aa-AMP) followed by transfer of the aminoacyl moiety to the tRNA (**Figure 1A**). Coupling of non-cognate aa-tRNA pairs can jeopardize translational fidelity, and this potentially damaging scenario is prevented by hydrolysis of misaminoacylated tRNA (post-transfer editing) at a separate aaRS editing domain. Some aaRS may also hydrolyze non-cognate aminoacyl-adenylate (pre-transfer editing) within the synthetic site [6]. Isoleucyl-tRNA synthetase (IleRS) decodes isoleucine codons. Misaminoacylation of the amino acids that closely mimic isoleucine, such as valine and non-proteinogenic norvaline, is prevented by IleRS pre- and post-transfer editing (**Figure 1A**) [7, 8]. IleRSs group in two distinct clades that differ in the structure of the anticodon binding domain, unique sequence elements of the catalytic site and susceptibility to the natural antibiotic mupirocin [8, 9]. The first clade (IleRS1) comprises most of bacterial as well as all eukaryote mitochondrial IleRSs. The members of the second clade (IleRS2), showing a significantly lower susceptibility to competitive inhibition by mupirocin [10], are found in some bacteria and in the eukaryotic cytoplasm. In general, bacteria rely on either *ileS1* or *ileS2* for housekeeping functionality. Interestingly, the simultaneous presence of both genomic *ileSs* in the same organism, besides mupirocin-producing *Pseudomonas fluorescens* NCIMB 10586 strain [11], appears to be unique to family *Bacillaceae* [12].

**Figure 1.**
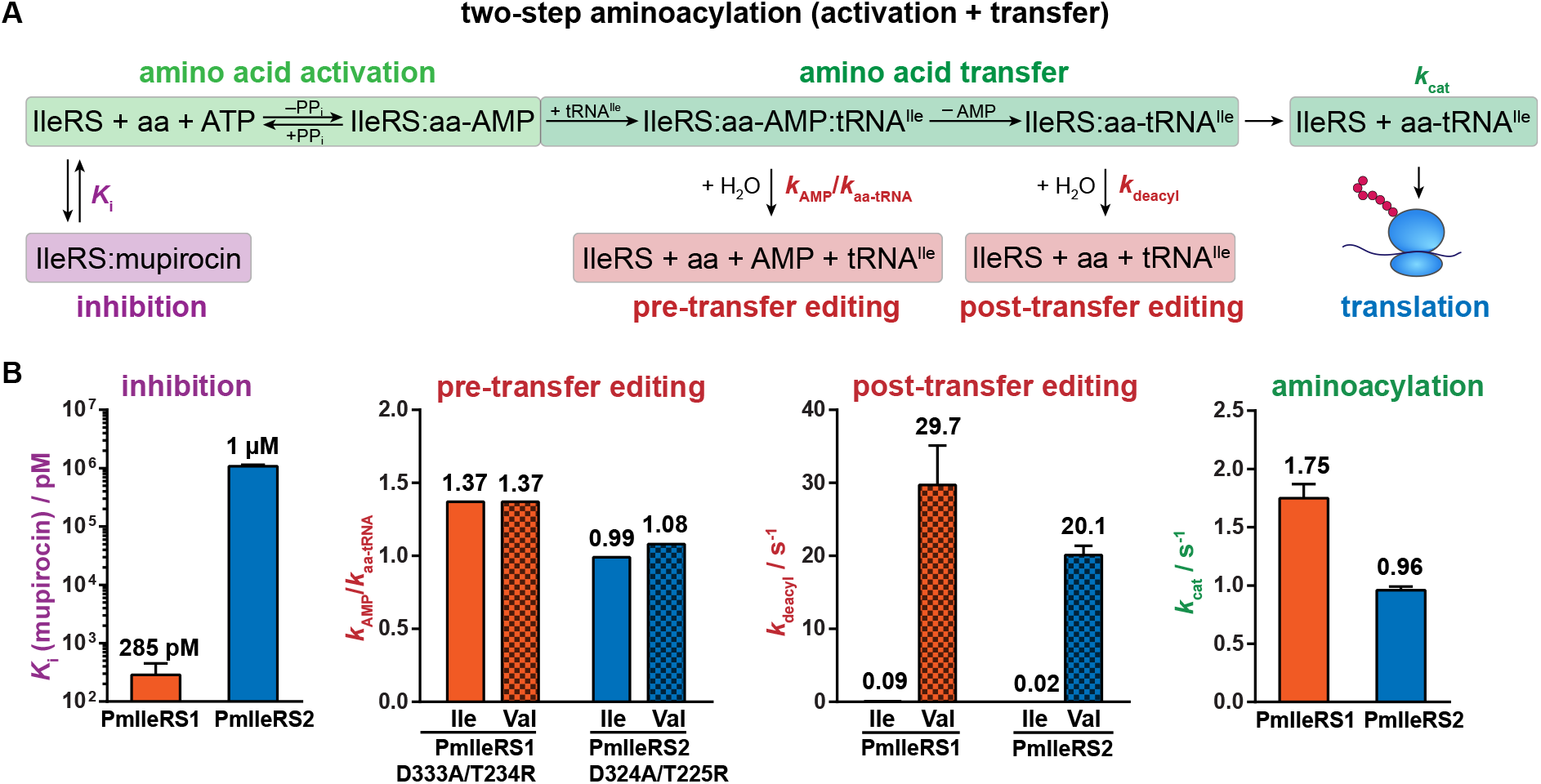
Schematic representation of IleRS aminoacylation and editing reactions with the corresponding key biochemical results. A) A two-step synthetic pathway (green) leads to synthesis of Ile-tRNA^Ile^, represented by aminoacylation turnover, *k*_cat_. IleRS inhibition by antibiotic mupirocin is described by inhibition constant, *K*_i_ (purple). The editing pathways (red) comprise i) tRNA-dependent pretransfer editing, calculated as the ratio of the steady-state rate constants for ATP consumption and aa-tRNA^Ile^ synthesis in the presence of cognate or non-cognate (Val) amino acid catalyzed by the post-transfer editing-defective IleRS variant (ratio higher than one indicates editing) and ii) post-transfer editing, represented by the single-turnover rate constant for aa-tRNA^Ile^ hydrolysis, *k*_deacyl_. B) Kinetic data unveil that PmIleRS2, relative to PmIleRS1, has a higher *K*_i_ for mupirocin (**Figure 3**), the same post-transfer editing rate (**Supplementary Table S3**) but lower aminoacylation *k*_cat_ (**Table 2**). Both PmIleRSs lack pre-transfer editing (**Supplementary Table S2**). The same editing results were obtained also with non-cognate Nva. Amino acid activation parameters are given in **Table 1**.

Here, we used comprehensive bioinformatics, kinetics, and mass spectrometry analyses to understand the reasons why these organisms maintain two *ileS* genes. Using *Priestia* (*Bacillus*) *megaterium* [13] as a model organism, we found that each paralogue provides the advantage of a distinct relevance. PmIleRS2 allows growth on mupirocin, while PmIleRS1 enables faster translation. Our data suggests that Bacilli, as generally fast-growing organisms, could not compromise with the slow catalytic turnover of PmIleRS2. Therefore, they rely on the IleRS1 for fast and efficient growth and benefit from the antibiotic resistance acquired *via* the second gene.

## Results

### Phylogenetic analysis suggests ileS1 gene is mandatory in Bacilli

Bacterial IleRS group into two distinct clades, IleRS1 and IleRS2, the latter carrying mupirocin resistance [8, 9]. The two enzymes share about 30 % identity (50-60 % similarity) [14], and each can stand as a sole housekeeping gene in bacteria. Yet, we have recently [8] noted a small bacterial niche within the family *Bacillaceae* in which the organisms carry both *ileS* genes in the same genome (**Figure 2**, purple clades). To explore that further, we performed comprehensive phylogenetic analysis of bacterial IleRSs. Surprisingly, we found that species of the family *Bacillaceae* always have *ileS1* while several species also have *ileS2* (i.e. *Priestia (Bacillus) megaterium* [13]). The rationale for acquiring the resistance-carrying *ileS2* is clear. However, what is not clear is whether *ileS2* can serve as the sole housekeeping gene in Bacilli as it does in other species like Actinobacteria and Chlamydia (**Figure 2**, blue clades). Indeed, a detailed public database search did not reveal a single species of the family *Bacillaceae* lacking *ileS1.* This indicates that IleRS1 is under strong positive selection in Bacilli.

**Figure 2.**
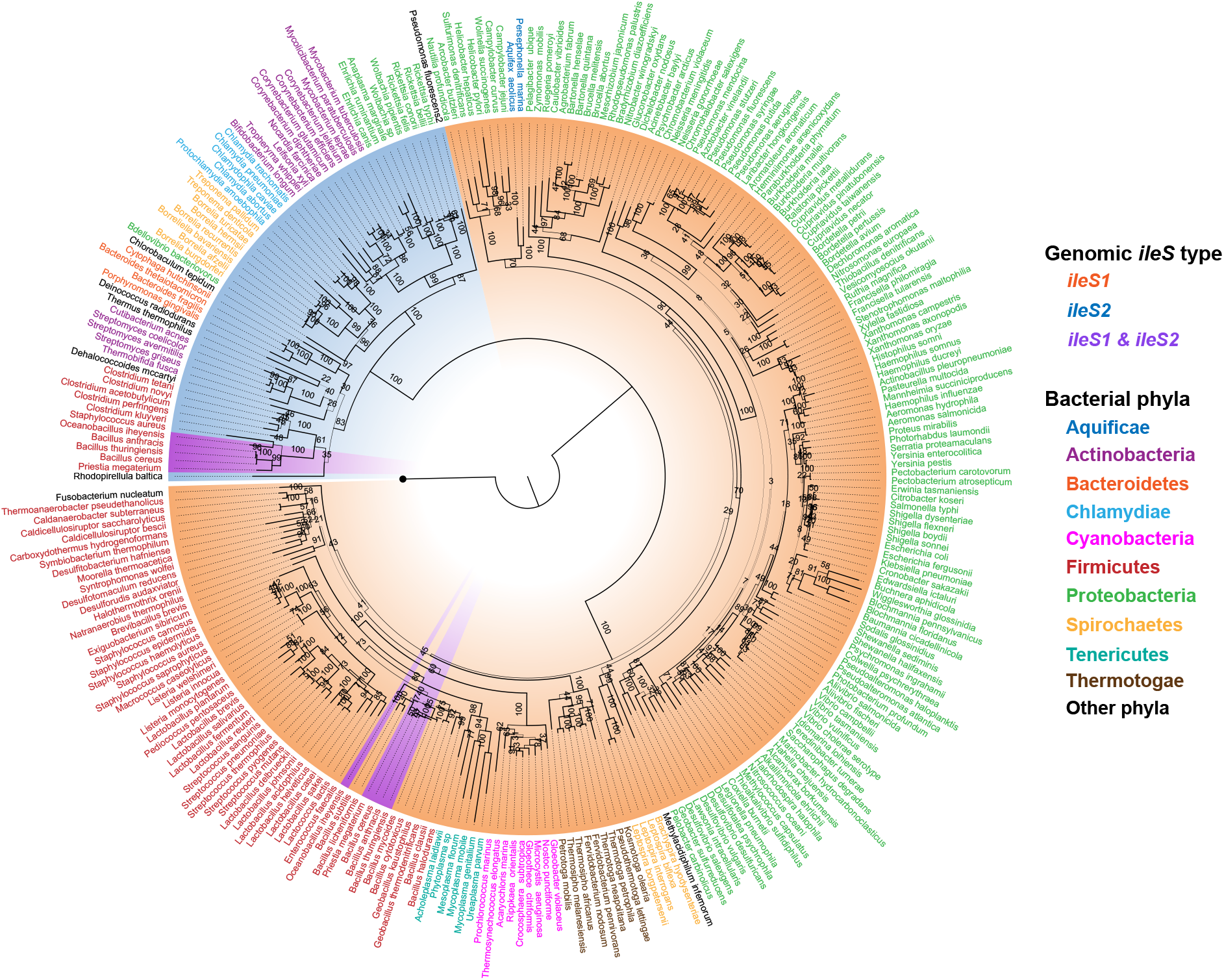
*ileS* distribution throughout bacterial kingdom. Phylogenetic tree was constructed using maximum likelihood method RAxML after multiple sequence alignment of 278 bacterial IleRS (218 *ileS1,* 60 *ileS2*). The tree was rooted using *E. coli* ValRS (black circle). The line width of each branch is scaled according to the bootstrap support value, and numbers along the nodes represent the percentage of nodes reproducibility in 1000 bootstrap replicates. Species names are colored by phylum. Nodes are highlighted according to the genomic *ileS* distribution. The species having both *ileS1* and *ileS2* (highlighted purple) belong exclusively to the family *Bacillaceae*. Exception is *Staphylococcus aureus* harboring plasmid copy of *ileS2*.

### PmIleRS2 binds mupirocin with 10^4^-fold lower affinity relative to PmIleRS1

To address a rationale for the mandatory presence of *ileS1,* we chose *P. megaterium* as a model organism and its two IleRSs as model enzymes (31 % identity). Homologous overexpression in *P. megaterium* and heterologous in *E. coli* yielded the enzymes with the same kinetic competence. Hence, the enzymes were predominately overexpressed in *E. coli* and purified using Ni-NTA. We started *in vitro* characterization by testing mupirocin inhibition (**Figure 1**). Analogously to stand-alone IleRS2s [8, 15], PmIleRS2 displayed a significant resistance to the antibiotic with a moderately high *K*_i_ (1 μM) measured at amino acid activation, the first step of the two-step aminoacylation reaction (**Figure 3A**). In contrast, PmIleRS1 was readily inhibited by nanomolar mupirocin and featured a more complex character with non-linear time courses characteristic for slow-binding inhibition (**Figure 3B left**). The *K*_i_ obtained for PmIleRS1 (**Figure 3B right**) is comparable with the *K*_i_ for *Staphylococcus aureus* IleRS1 [16]. Thus, PmIleRS1 and PmIleRS2 display a 10^4^-fold difference in sensitivity to mupirocin in accordance with the prediction that PmIleRS2 provides mupirocin resistance to *P. megaterium.*

**Figure 3.**
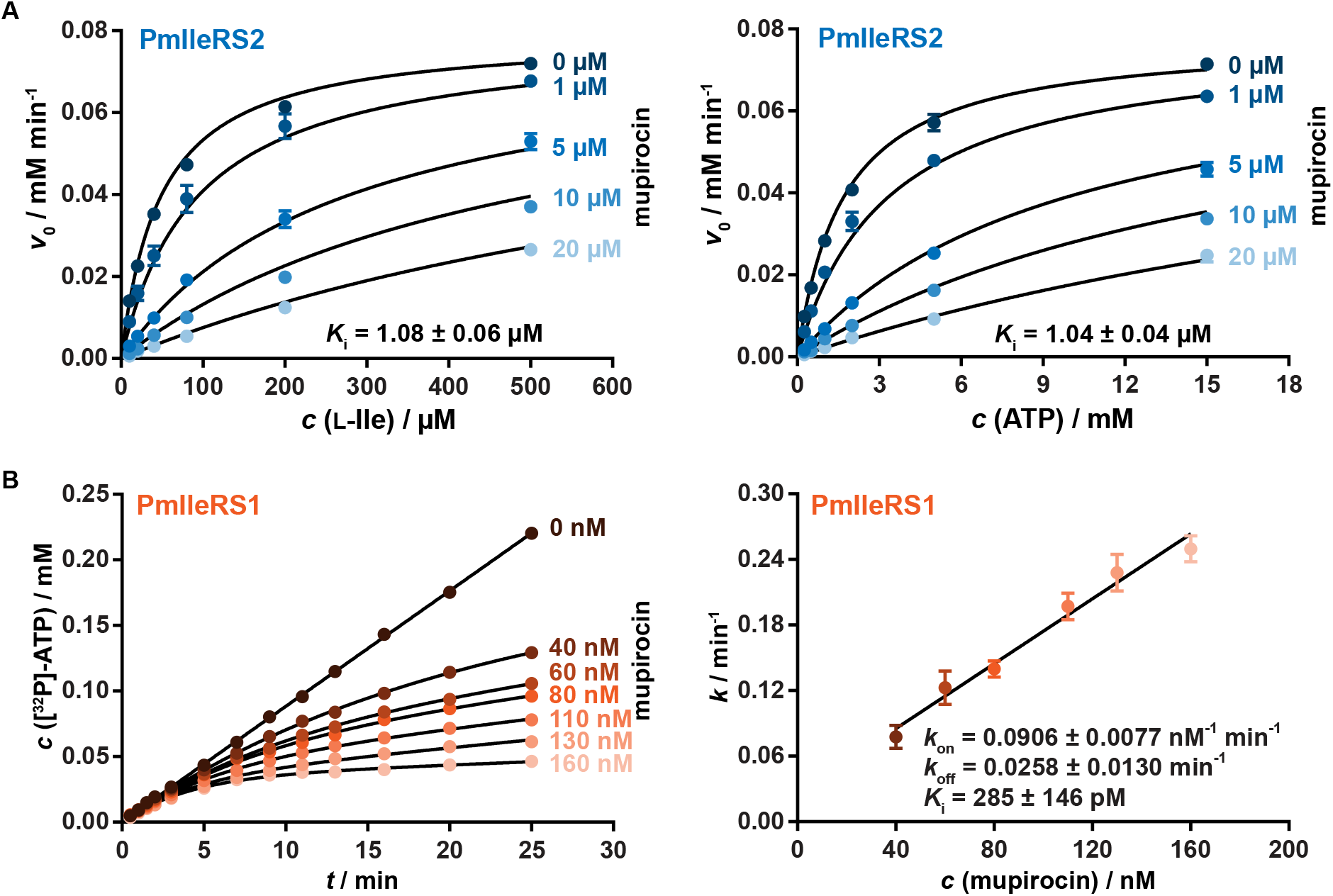
Inhibition of PmIleRS1 and PmIleRS2 by mupirocin. Values represent the average value ± SEM of at least three independent experiments and error bars correspond to SEM. A) PmIleRS2 displayed a classic fast-on/fast-off competitive inhibition in Ile activation. The figure depicts plots of initial reaction rates vs substrate concentration (left Ile, right ATP) in the presence of increasing concentrations of mupirocin. Data were fitted to classic competitive inhibition equation to obtain the inhibition constant (*K*_i_). B) PmIleRS1 displayed a slow, tight-binding competitive inhibition. The time-courses (left) were fitted to a slow-binding equation and the apparent rate constant *k* was replotted vs mupirocin concentration to reach the *k*_on_ and *k*_off_ describing mupirocin binding. Inhibition constant (*K*_i_) was calculated from the ratio of *k*_off_ and *k*_on_ (for details see Supplementary Materials and methods).

### PmIleRS1 is two-fold faster in Ile-tRNA^Ile^ synthesis than PmIleRS2

PmIleRS2 insensitivity to mupirocin could have evolved through a trade-off with the enzyme’s efficiency and/or accuracy. To explore this possibility, we first compared the PmIleRS1 and PmIleRS2 catalytic competences. We found that PmIleRS1 is indeed 10-fold more efficient in isoleucine activation (measured as the *k*_cat_/*K*_M_) relative to PmIleRS2 (**Table 1**). Along the same line, PmIleRS1 appeared superior to PmIleRS2 in the two-step aminoacylation by showing a two-fold higher catalytic turnover (*k*_cat_, **Table 2, Supplementary Figure S1**). However, at the level of *k*_cat_/*K*_M_ PmIleRS1 equals PmIleRS2, due to the opposing *K*_M_ (Ile) trends in overall aminoacylation (**Table 2**) and the isolated activation step (**Table 1**). Although the origin of this effect is not completely clear, an increase in the aminoacylation *K*_M_ (Ile) for PmIleRS1 can be attributed to the modulation of Ile recognition by tRNA (**Supplementary Table S1**) [7, 17]. Nevertheless, given the cellular concentration of Ile (~ 300 μM) [18], PmIleRS1 and PmIleRS2 may operate at near saturating conditions *in vivo*. Thus, the higher aminoacylation turnover (*k*_cat_) of PmIleRS1 may be relevant in the cell.

**Table 1.**
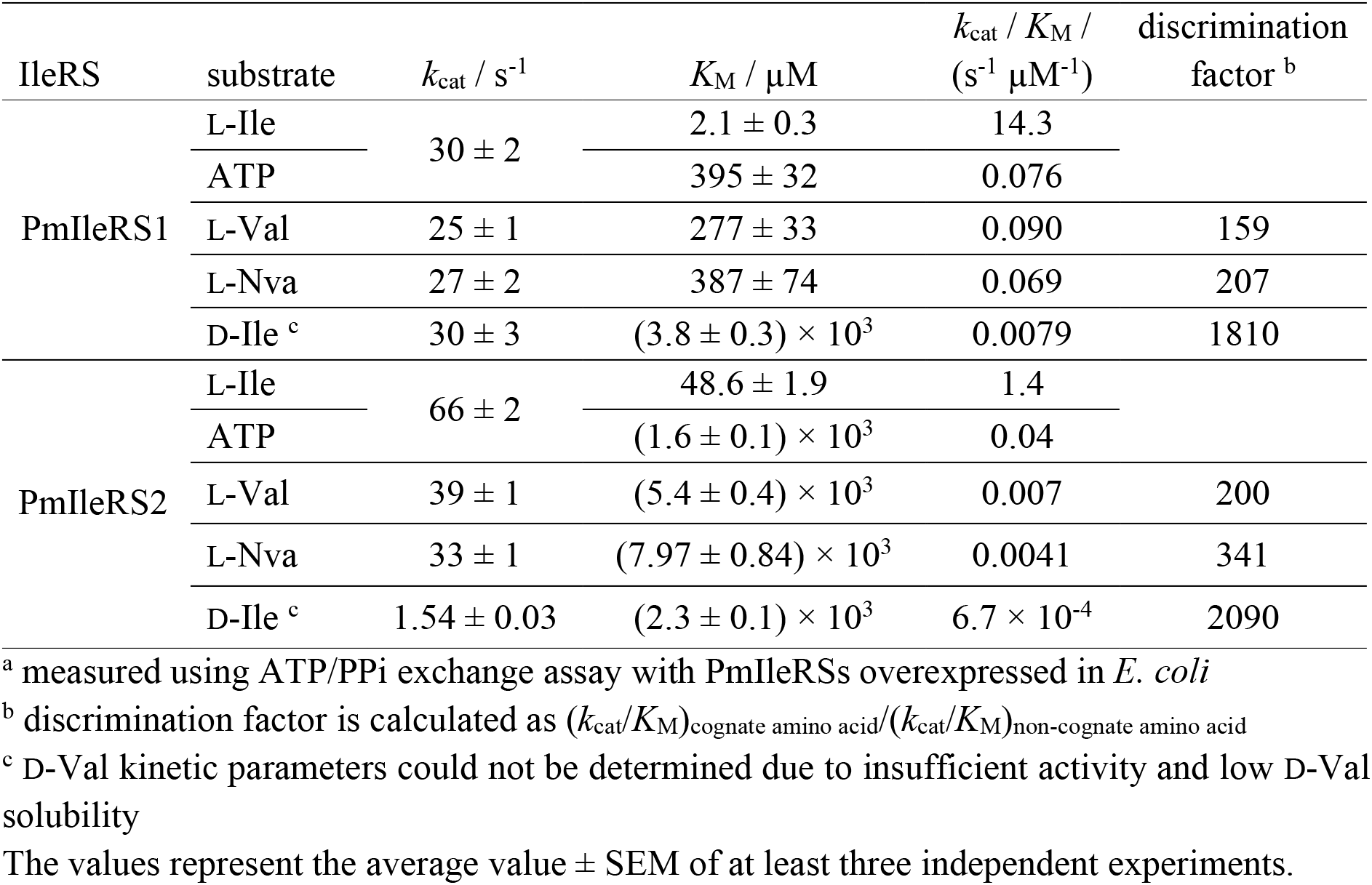
Steady state amino acid activation^*a*^.

### PmIleRS1 maintains ribosomal translation better than PmIleRS2

To what extent might a 2-fold difference in aminoacylation turnover influence protein translation? This question was addressed using a cell-free coupled transcription-translation system (PURExpress, NEB) deprived of *E. coli* IleRS. PmIleRS1 or PmIleRS2 were added at the same concentrations and synthesis of a dihydrofolate reductase (DHFR) reporter enzyme containing 12 Ile residues was monitored by incorporation of [^35^S]-methionine. Under conditions where protein translation was limited by Ile-tRNA^Ile^ synthesis, PmIleRS1 accumulated more DHFR compared to PmIleRS2, especially at prolonged translation (> 2 hours, **Figure 4**). Importantly, under these conditions stability and activity of PmIleRS2 was not affected (**Supplementary Figure S2**). Thus, the cell-free data endorse *in vitro* kinetic analysis by showing a noticeable superiority of PmIleRS1 in translation.

**Figure 4.**
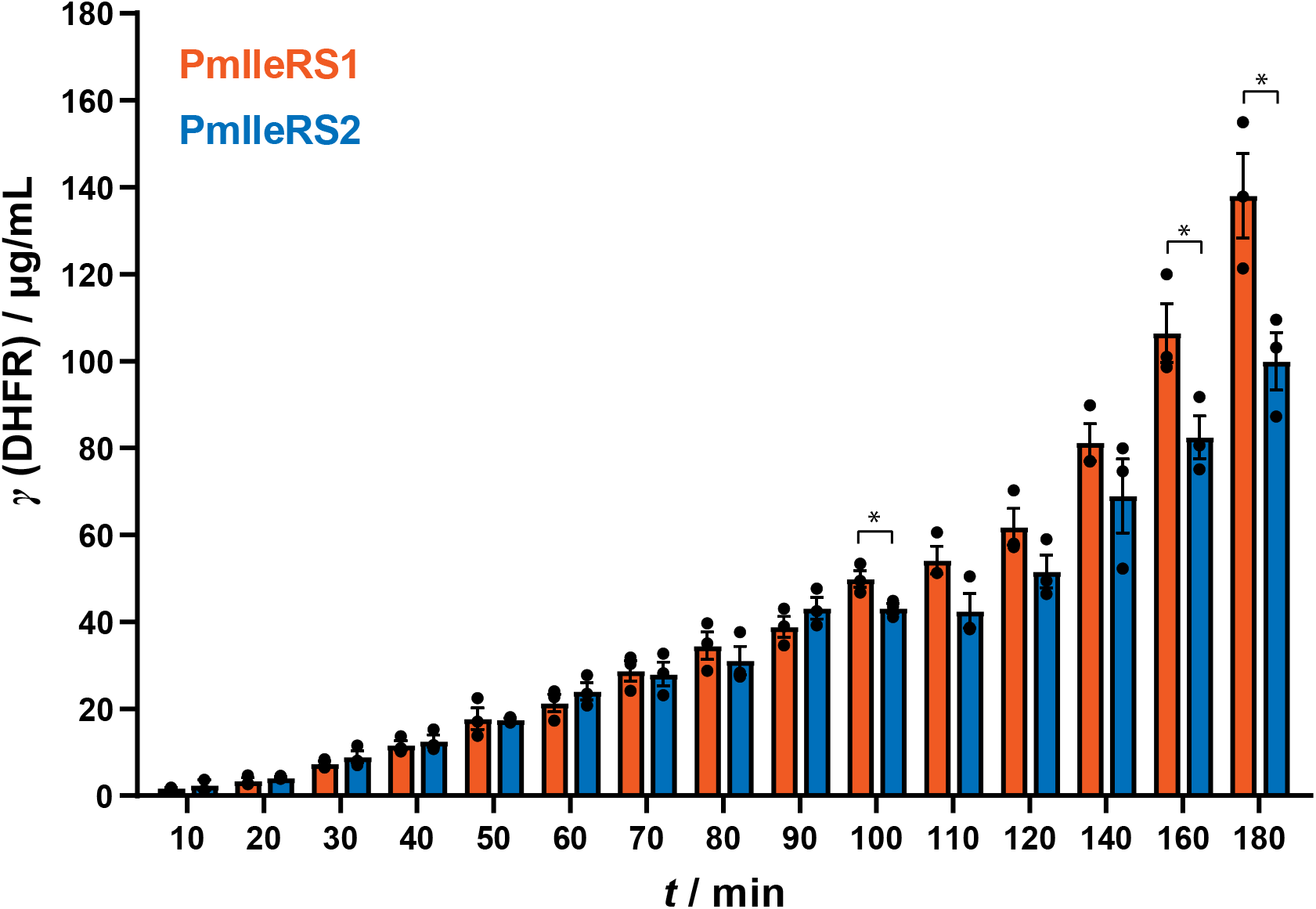
*In vitro* translation of dihydrofolate reductase using PmIleRSs. The values represent the average value ± SEM of three independent experiment and error bars correspond to SEM. Statistically significant difference in DHFR amount was calculated using Student’s t-test and asterisk (*) indicates statistical difference with p < 0.05.

### Discrimination against non-cognate amino acids is conserved between PmIleRSs

The next question was whether the PmIleRS2 fidelity is compromised due to the increased insensitivity to mupirocin. Therefore, activation of the near-cognate, L-Val and L-Nva, and also D-Ile was tested, and the discrimination factors (*D*) calculated (**Table 1**). Low *D* implies frequent misactivation and, based on the anticipated tolerable translational error [19], *D* < 3300 is taken as an indication for editing. The data show that, analogously to other IleRSs [7, 8], PmIleRS1 and PmIleRS2 discriminate against L-Val equally well as against L-Nva. Discrimination is based on a ~150-fold higher *K*_M_ for L-Val or L-Nva relative to the cognate Ile, while the *k*_cat_ remains almost the same (**Table 1**). Both enzymes also discriminate similarly against D-Ile, pointing that the existence of the two enzymes is not related to the possible use of D-cognate amino acids in peptide wall synthesis, as in the case of *B. subtilis* TyrRS [20]. Thus, in contrast to the catalytic turnover which appears to be compromised in PmIleRS2, the enzymes do not differ in initial amino acid selectivity.

### Both enzymes feature post-transfer editing and lack pre-transfer editing

Initial amino acid selectivity and editing jointly modulate IleRS accuracy. So, we next questioned whether PmIleRS1 and PmIleRS2 differ in their post-transfer editing capacity (**Figure 1**). We prepared Val- and Nva-[^32^P]-tRNA^Ile^ and mixed them with an excess of enzyme (see Supplementary Materials and methods) to measure the first-order rate constants describing the hydrolysis of misacylated tRNAs (*k*_deacyl_). The obtained rates were similar (**Supplementary Table S2**) demonstrating that PmIleRS1 and PmIleRS2 resemble each other in post-transfer editing. The rates were about 10^4^-fold faster than the non-enzymatic hydrolysis (**Supplementary Table S2**) showing that PmIleRSs, like other IleRSs [7, 8], feature rapid posttransfer editing.

Besides post-transfer editing, error correction at the level of misaminoacyl-AMPs (pre-transfer editing, **Figure 1**) was previously reported for *E. coli* IleRS [21, 22]. This editing pathway can be separately measured using mutants disabled in post-transfer editing [23]. To explore whether the *P. megaterium* paralogues also feature this editing step, the post-transfer editing inactive variants, PmIleRS1 D333A/T234R and PmIleRS2 D324A/T225R (**Supplementary Table S2**), were tested in parallel steady-state assays that follow the formation of aa-tRNA^Ile^ (*k*_aa-tRNA_) and the consumption of ATP (measured as AMP formation, *k*_AMP_) [24]. The mutants consumed stoichiometric amount of ATP per synthesized aa-tRNA (*k*_AMP_/*k*_aa-tRNA_ ratios for Nva and Val close to 1; **Supplementary Table S3**), demonstrating that both PmIleRS paralogues lack pretransfer editing. As expected, the WT enzymes exhibit *k*_AMP_/*k*_aa-tRNA_ ratios for Val and Nva substantially higher than 1 (**Supplementary Table S3**) in agreement with strong post-transfer editing. In summary, PmIleRS paralogues exhibit no difference in the accuracy of aminoacylation. The main distinction between the paralogues remains the aminoacylation turnover, which is lower in PmIleRS2 as a possible trade-off with mupirocin resistance.

### Expression of PmIleRS2 is highly upregulated by mupirocin

Does the rationale for PmIleRS1 requirement lie, besides its kinetic superiority, also in its regulation? To address this question, we performed bioinformatic analyses that indicated a housekeeping mode of regulation for *ileS1* with its promoter being recognized by RNA polymerase containing σ^70^. In contrast, the *ileS2* expression seems to be regulated by CodY repressor (**Figure 5A**). CodY is a GTP-sensing repressor protein involved in stringent response in Gram-positive bacteria. Its repression is relived at low GTP concentration [25]. We also found conserved T-Box riboswitch structural elements preceding both *ileS* ORFs (**Supplementary Figure S3B**). This regulation mechanism, in Gram-positive bacteria, senses the aminoacylation state of the tRNA and dictates either read-through or termination of transcription via tRNA:mRNA interactions [26]. Furthermore, detained analysis revealed that *ileS1* location in the genome is well conserved throughout *Bacillaceae* species, unlike *ileS2* location which is only conserved in the small subpopulation of Bacilli species (**Supplementary Figure 4**).

**Figure 5.**
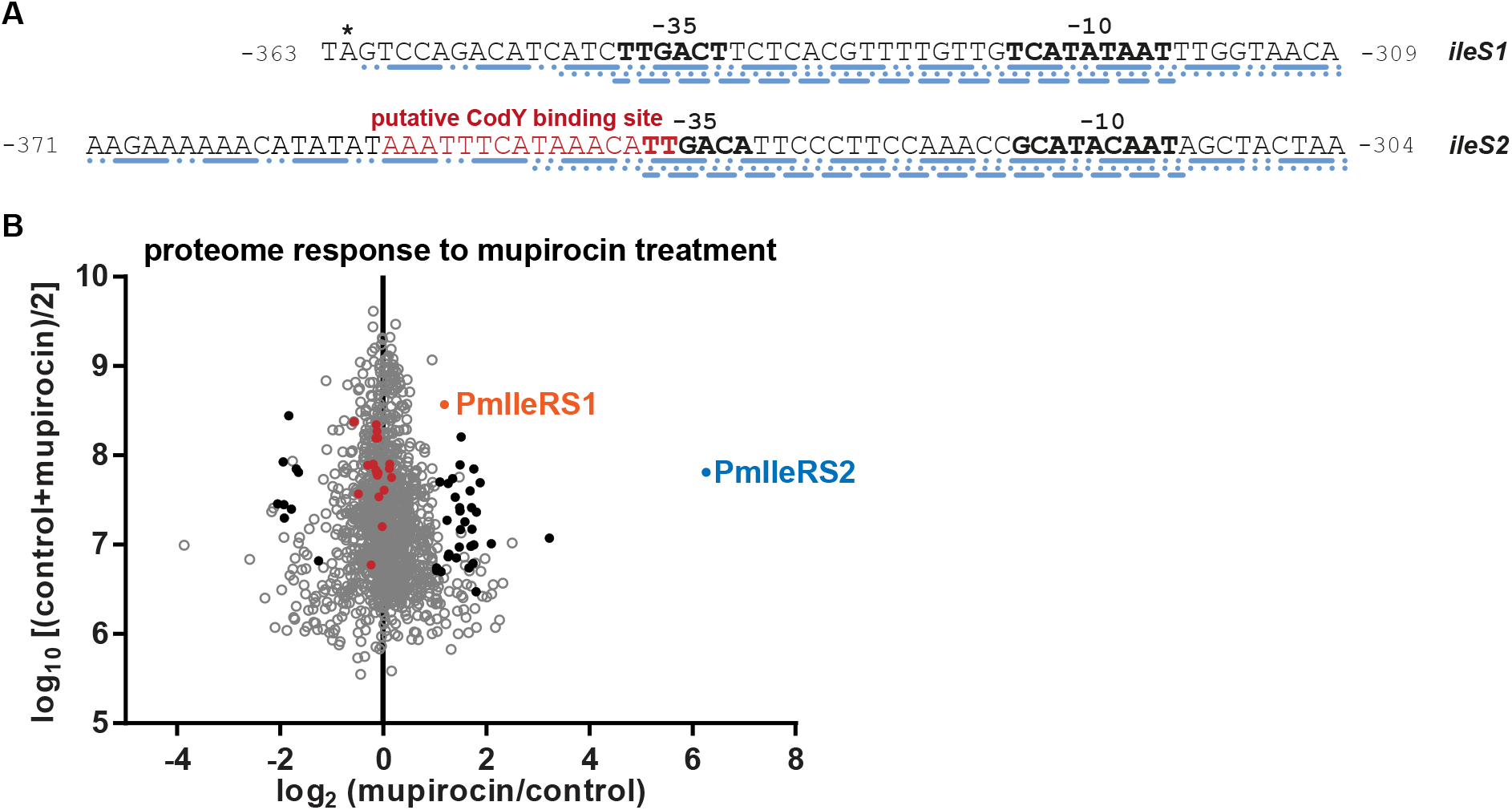
Putative transcriptional regulation and expression profiles of *P. megaterium* IleRSs. A) Upstream region of *ileS* genes analyzed by BacPP 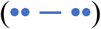, PePPER 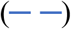 and FruitFly 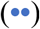. Numbering starts from the first nucleotide of the ORF (not shown). T-box regulatory sequences (not shown, **Supplementary Figure S3A)** are inserted between the first nucleotide of the ORF and a putative −10 promoter element; asterisk (*) denotes the stop codon of the upstream gene. SoftBerry analysis of *P. megaterium* putative −35 and −10 promoter elements (bold) indicate that *rpoD16* and *rpoD19* transcription factors (both part of σ^70^ subunit) regulate *ileS1* while *ileS2* is regulated by stress repressors. A putative CodY binding site is localized within −35 promoter region. B) Mass spectrometry proteome analysis revealed 70-fold upregulation of PmIleRS2 in the presence of mupirocin. Filled black circles represent proteins with more than 2-fold statistically significant change in expression including also PmIleRS1 (two sided, two sample Student’s t-test with Benjamini-Hochberg correction for multiple hypothesis testing, FDR=0.05). The red circles mark other aaRSs.

The anticipated PmIleRS1 and PmIleRS2 expression profiles were tested by mass spectrometry (MS). Without mupirocin (the control sample), PmIleRS1 was readily detectable, while the PmIleRS2 peptides were close to the detection limit. Addition of mupirocin upregulated PmIleRS2 expression by approximately 70-fold, PmIleRS1 by 2-fold, and did not affect expression of any other aaRS (**Figure 5B**). What is the mechanism by which mupirocin induces *ileS* genes expression? The most plausible model includes PmIleRS1 inhibition and subsequent accumulation of non-aminoacylated tRNA^Ile^. The tRNA/Ile-tRNA^Ile^ ratio may transmit the signal (of inactive PmIleRS1) by promoting i) the stringent response [27] which in turn stimulates dissociation of the repressor and ii) the read-through T-box conformation that stimulates transcription. Thus, *ileS2* expression may arise from the combined action of these two pathways (**Supplementary Figure S3B**). Less intuitive is the effect exerted on PmIleRS1 (please note the magnitude of the effect is much smaller). However, one has to keep in mind that *ileS1* expression from the constitutive promotor is also tuned by the T-box (**Supplementary Figure S3B)**, and thus autoregulated by IleRS1 inhibition by mupirocin. SDS-PAGE fractionation followed by in-gel digestion improved sequence coverage and allowed for a better estimate of the relative abundance of PmIleRS1 and PmIleRS2 (**Supplementary Figure S5**). These data are consistent with PmIleRS1 being constitutively expressed while PmIleRS2 is induced by antibiotic stress to the slightly lower level than PmIleRS1 in the absence of antibiotic.

### PmIleRS2 can provide P. megaterium with housekeeping functionality and mupirocin resistance

Our bioinformatic analysis revealed that *ileS2* never stands alone in Bacilli. So, a question emerged as to whether PmIleRS2, and specifically under its native inducible promoter, could serve as the sole housekeeping enzyme in Bacilli. To investigate this, we constructed both *P. megaterium* strains deleted for *ileS1* or *ileS2* (**Supplementary Figure 6**). As expected, the *ΔileS2* construction was straightforward and led to mupirocin sensitivity. The latter was recovered by expression of PmIleRS2 under its native promoter, from a plasmid (**Figure 6B, Supplementary Figure S7**). Loss of *ileS2* did not influence the growth in the absence of mupirocin, strongly arguing this gene is related only to the resistance (**Supplementary Figure S8**).

**Figure 6.**
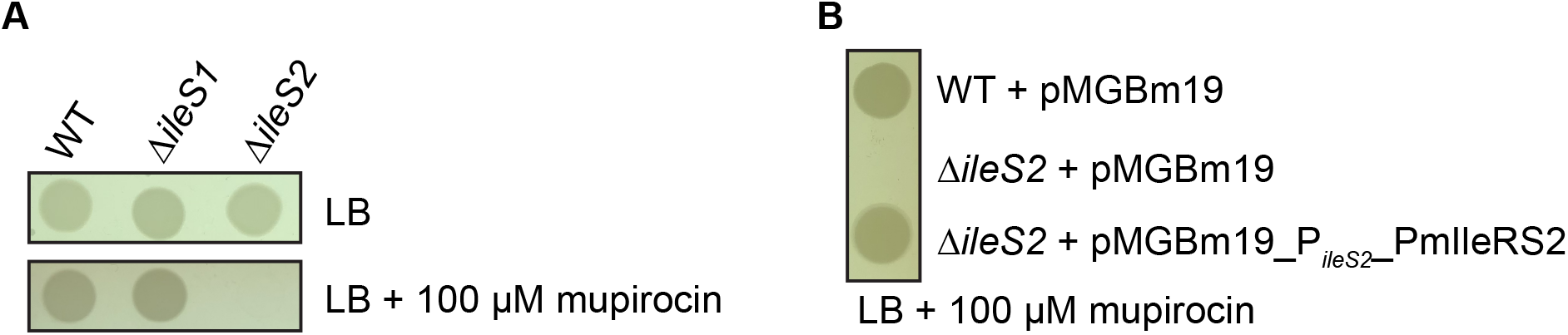
Phenotypic characterization of the *P. megaterium* knockout strains. A) Both *ΔileS1 and ΔileS2* were viable in the absence of mupirocin. 100 μM mupirocin arrested growth of *ΔileS2.* B) *ΔileS2* mupirocin-sensitive phenotype was complemented by *ileS2* expressed from its native promoter from the plasmid pMGBm19_P_*ileS2*__PmIleRS2. PmIleRS2 expression was confirmed by Western blot (**Supplementary Figure S7**).

In sharp contrast, the *ΔileS1* strain could be selected only when the cells were grown in the presence of mupirocin. The constructed strain was nevertheless viable without mupirocin indicating that expression of PmIleRS2 is constitutive in the absence of *ileS1* (**Figure 6A)**. However, the whole-genome sequencing of the *ΔileS1* strain revealed that the strain has lost two out of six plasmids (carrying approximately 200 genes) naturally found in the WT strain. This may arise because deletion of the *ileS1* gene *per se* comes at the expense of genomic stability or due to the selection in the presence of mupirocin. Whatever the reason is, our data demonstrate that PmIleRS2, under its native promoter can support housekeeping functionality along with antibiotic resistance. We propose this solution is extremely rare (if it exists) in Bacilli due to the speed-resistance trade-off in IleRS2.

## Discussion

Simultaneous presence of two paralogues in a single cell can reflect a need for: i) distinct enzymatic features, ii) differential regulation, or iii) both [20, 28, 29]. We investigated two IleRS paralogues from *P. megaterium,* each belonging to a distinct clade of bacterial IleRS proteins (dubbed IleRS1 and IleRS2 [8, 9]). PmIleRS2 displayed insensitivity to the antibiotic mupirocin, with the *Ki* in the micromolar range, whereas PmIleRS1 was sensitive to nanomolar concentrations of the antibiotic (**Figure 3**). Consequently, PmIleRS2, as it is the case for other characterized IleRS2 enzymes [8, 15, 30], supports antibiotic resistance (**Figure 6**), a feature that likely influence the horizontal gene transfer events observed in *ileS2* [8]. Interestingly, although *ileS2* can stand alone in a number of bacteria, in Bacilli, *ileS2* is present only as a second gene (**Figure 2**). This opens the intriguing question of whether IleRS2-only organisms experience fitness trade-offs associated with the acquired mupirocin resistance [31]. We rationalized that either aminoacylation fidelity, known to ensure cell viability [7, 32], or aminoacylation speed could be compromised in IleRS2. Detailed kinetic analysis demonstrated that PmIleRS2 is indistinguishable from PmIleRS1 (and other previously tested IleRS1 and IleRS2 enzymes [7, 8]) in terms of initial substrate selectivity (**Table 1**) or post-transfer editing **(Supplementary Table S2**). Interestingly, PmIleRS2, similar to some other IleRS2 proteins [8], lacks tRNA-dependent pre-transfer editing within the synthetic site (**Supplementary Table S3**). However, we found the same for PmIleRS1 (**Supplementary Table S3**). Taken together, the evolutionary pressure for IleRS1 in Bacilli is apparently not related to aminoacylation fidelity. However, we discovered that the mupirocin resistant PmIleRS2 has two-fold slower aminoacylation turnover (*k*_cat_) (**Table 2**), which translates into a noticeably slower cell-free translation (**Figure 4**), suggesting that mupirocin-resistance may come on account of the translational rate. Could this govern the universal presence of IleRS1 in Bacilli, and even more broadly? Limited kinetic data obtained here and reported previously on related systems show that IleRS1s are indeed faster in aminoacylation than IleRS2s (**Table 2**, [8, 33]). Also, we collected data on the minimum doubling times of 207 bacteria from the literature and compared this with IleRS distribution (**Figure 7**). These data revealed that median of the doubling time of bacteria comprising IleRS1 is at least 2-fold lower (i.e. faster growth) than of bacteria relying exclusively on IleRS2. Thus, it appears that IleRS1 is required by faster-growing bacteria to maintain fast translation (**Supplementary Figure S9**). In summary, our data provide evidence that IleRS1 in Bacilli and other fast-growing bacteria is maintained due to selective pressure for fast translation. In contrast, slow-growing bacteria may trade-off translational speed for the higher mupirocin resistance and rely on IleRS2.

**Figure 7.**
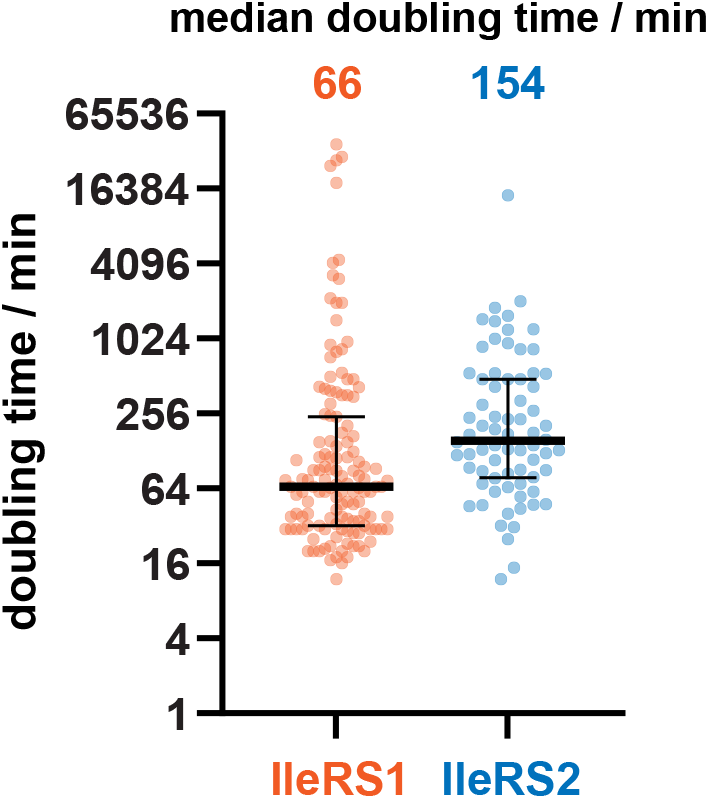
Relationship between IleRS type and bacterial doubling time. Each dot represents one bacterial species colored according to the genomic IleRS type (n (IleRS1) = 135, n (IleRS2) = 72). Doubling times were compiled from the literature (if multiple conditions were reported the minimal doubling time was selected) and the median doubling times calculated.

**Table 2.** Steady state two-step aminoacylation^*a,b*^.

| IleRS | substrate | $k_{\text{cat}} / \text{s}^{-1}$ | $K_{\text{M}} / \mu\text{M}$ | $k_{\text{cat}} / K_{\text{M}} / (\text{s}^{-1} \mu\text{M}^{-1})$ |
| --- | --- | --- | --- | --- |
| PmIleRS1 | L-Ile | $1.75 \pm 0.12$ | $43.2 \pm 3.3$ | 0.04 |
| | ATP | | $246.0 \pm 19.8$ | 0.01 |
| | tRNA<sup>Ile</sup> | | $2.03 \pm 0.15$ | 0.86 |
| PmIleRS2 | L-Ile | $0.96 \pm 0.03$ | $6.2 \pm 0.8$ | 0.15 |
| | ATP | | $76.4 \pm 1.4$ | 0.01 |
| | tRNA<sup>Ile</sup> | | $3.7 \pm 0.3$ | 0.26 |

<sup>a</sup> measured using PmIleRSs overexpressed in *P. megaterium*
<sup>b</sup> the progress curves displayed biphasic nature. The fast phase was used to determine the kinetic parameters (**Supplementary Figure 2**).
The values represent the average value ± SEM of at least three independent experiments.

## Materials and methods

### Bacterial strains, growth conditions and cloning

All cloning was performed in *E. coli* as described [34] or following manufacturer’s protocol. All plasmids, PCR products and strains were verified by sequencing. Whole-genome sequencing was performed on Illumina platform (2 x 150 bp, paired ends) (Macrogen). All plasmids are listed in **Supplementary Table S4**, primers in **Supplementary Table S5**, and plasmid construction details in **Supplementary Table S6**. *P. megaterium* (DSMZ, strain DSM-32) was routinely grown at 30 °C in LB medium. Knockout strains were prepared as described [35].

### Production of IleRS enzymes and tRNA^Ile^

*P. megaterium* enzymes (WT and variants) were cloned (**Supplementary Tables S4, S5 and S6**) as a N-terminal His_6_-tagged proteins and overexpressed from pET28 in *E. coli* BL21(DE3) at 30 °C (PmIleRS1) or 15 °C (PmIleRS2) using 0.25 mM IPTG. Enzymes were purified by standard Ni^2+^-NTA affinity chromatography [33]. *P. megaterium* tRNA^Ile^_GAU_ gene was cloned under T7 promoter and overexpressed from pET3a in *E. coli* BL21(DE3) at 30 °C using 1 mM IPTG [33]. Protein expression in *P. megaterium* was carried out from pP_T7_ (for PmIleRS1) and pMGBm19 (for PmIleRS2) plasmids at 30 °C using 0.5 % xylose [36].

### Kinetic assays

All assays were performed at 30 °C essentially as described [8, 33]. Slow, tight binding mupirocin inhibition of PmIleRS1 was assayed as described by Morrison and Walsh [37]. For details of all kinetic assays see Supplementary Materials and methods.

*In vitro* coupled transcription-translation was performed at 37 °C using the PURExpress ΔIleRS Kit (NEB) following manufacturer’s protocol supplemented with [^35^S]-methionine (EasyTag, Perkin Elmer) and 20 U murine RNase inhibitor (NEB). PmIleRS enzymes were present at 4 nM. Quantification of reporter protein (dihydrofolate reductase) was performed following manufacturer’s protocol.

### Bioinformatics and phylogenetic analysis

Phylogenetic tree was constructed as described [8]. Putative promoter elements were analyzed using BacPP [38], FruitFly [39] and PePPER [40] bioinformatics tools and the putative transcription factor binding sites were determined by BProm [41] using approximately 500 nucleotides upstream of the *ileS* ORF as a query.

### MS analysis

For the whole proteome analysis, tryptic digests were prepared as described [42]. The digests were analyzed on nanoLC-MS/MS system (EASY-nLC 1200 coupled to Q Exactive Plus mass spectrometer, ThermoFisher Scientific) equipped with EASY-Spray™ HPLC C18 Column (2 μm particle size, 75 μm / 250 mm). ProteomeDiscoverer 2.4 and MaxQuant 1.6.17 were used for data analysis, protein identification and label-free quantification. All samples were prepared and analyzed in triplicates.

## Data availability

The raw mass spectrometry data and analysis results have been deposited to the ProteomeXchange with identifier_________.

## Author contribution

**Vladimir Zanki**: investigation (lead), methodology (lead), validation (lead), visualization (lead), writing – original draft preparation (equal), writing – review & editing (equal) **Bartol Bozic**: investigation (supporting), validation (supporting), writing – review & editing (supporting)

**Marko Mocibob**: investigation (supporting), validation (supporting), writing – review & editing (supporting)

**Nenad Ban**: funding acquisition (equal), writing – review & editing (supporting) **Ita Gruic-Sovulj**: conceptualization (lead), funding acquisition (equal), project administration (lead), supervision (lead), writing – original draft preparation (equal), writing – review & editing (equal)

## Acknowledgments

We gratefully acknowledge Jeff Errington, Lianet Noda-Garcia, Aleksandra Marsavelski, Igor Zivkovic, Valentina Evic, Mario Kekez, Alojzije Brkic and Kian Bigović Vill for fruitful discussions regarding the experiments and critical reading of the manuscript, Igor Zivkovic and Maja Baraci for the technical support with the experiments, and Rebekka Biedendieck for providing us with all the plasmids for *B. megaterium.* I. G.-S. is grateful to Dan S. Tawfik for his highly positive spirit in science and beyond, and for numerous vivid discussions about the IleRS1 and IleRS2 evolution and function.

