## Supplementary Materials for "A pair of isoleucyl-tRNA synthetases in Bacilli fulfill complementary roles to keep fast translation and provide antibiotic resistance"

**Supplementary Tables:**

**Supplementary Table S1.** Steady state amino acid activation in the presence of $tRNA^{Ile}_{ox}$

**Supplementary Table S2.** Single-turnover deacylation

**Supplementary Table S3.** Formation of [$^{32}$P]-AMP and aa-[$^{32}$P]-$tRNA^{Ile}$ using WT and post-transfer editing defective variants

**Supplementary Table S4.** List of the plasmids used in this study

**Supplementary Table S5.** List of the oligonucleotides used in this study

**Supplementary Table S6.** Construction of the plasmids used in this study

**Supplementary Table S7.** Bacteria doubling time

**Supplementary Figures:**

**Supplementary Figure S1.** Biphasic nature of $BmtRNA^{Ile}$ aminoacylation profiles

**Supplementary Figure S2.** PmIleRSs stability and aminoacylation activity after prolonged preincubation

**Supplementary Figure S3.** Putative regulation of *ileS* gene expression in *Bacillaceae*

**Supplementary Figure S4.** Genomic location of *ileS* genes in *Bacillaceae*

**Supplementary Figure S5.** Relative abundance of PmIleRSs

**Supplementary Figure S6.** Schematic presentation of the *P. megaterium ΔileSx* strain design

**Supplementary Figure S7.** Western-blot analysis of PmIleRSs expression

**Supplementary Figure S8. Growth analysis of *P. megaterium* strains**

**Supplementary Figure S9**. Relationship between bacterial phyla and the doubling time

**Supplementary Materials and Methods**

**Supplementary References**

**Supplementary Table S1. Steady state amino acid activation in the presence of $tRNA^{Ile}_{ox}$ [a, b]**

| IleRS | substrate | $k_{cat}$ / $s^{-1}$ | $K_M$ / µM | $k_{cat}$ / $K_M$ / ($s^{-1}$ $µM^{-1}$) |
|---|---|---|---|---|
| PmIleRS1 | L-Ile | 24 ± 3 | 16.3 ± 0.4 | 1.47 |
| | ATP | | (2.3 ± 0.3) × $10^3$ | 0.01 |
| PmIleRS2 | L-Ile | 65 ± 4 | 69.3 ± 5.5 | 0.94 |
| | ATP | | (3.8 ± 0.2) × $10^3$ | 0.02 |

[a] measured using ATP/PPi exchange assay with PmIleRSs overexpressed in *E. coli*

[b] $tRNA^{Ile}_{ox}$ is the oxidized variant of $tRNA^{Ile}$ with vicinal diols at the terminal ribose oxidized and thus unable to attach the amino acid. $tRNA^{Ile}_{ox}$ was prepared as described [1]. $tRNA^{Ile}_{ox}$ was renatured before use and was present at 10 µM concentration in the reaction mixture.

The values represent the average value ± SEM of three independent experiments.

**Supplementary Table S2. Single-turnover deacylation** [a]

| | $\boldsymbol{k}_{\mathbf{deacyl}}$ **/ s$^{-1}$** | | |
|---|---|---|---|
| IleRS | Ile-[$^{32}$P]-tRNA$^{Ile}$ | Val-[$^{32}$P]-tRNA$^{Ile}$ | Nva-[$^{32}$P]-tRNA$^{Ile}$ |
| PmIleRS1 WT [b] | 0.086 ± 0.003 | 29.7 ± 5.4 | 34.0 ± 5.6 |
| PmIleRS2 WT | 0.020 ± 0.001 | 20.1 ± 1.3 | 27.4 ± 1.5 |
| PmIleRS1 D333A/T234R | 0.0015 ± 0.0001 | 0.0004 ± 0.0001 | 0.0016 ± 0.0001 |
| PmIleRS2 D324A/T225R | 0.0003 ± 0.0001 | 0.0005 ± 0.0001 | 0.0008 ± 0.0001 |
| No enzyme | 0.0003 ± 0.0001 | 0.0006 ± 0.0002 | 0.0008 ± 0.0001 |

[a] measured using PmIleRSs overexpressed in *E. coli*

[b] hydrolysis of both Val-[$^{32}$P]-tRNA$^{Ile}$ and Nva-[$^{32}$P]-tRNA$^{Ile}$ displayed biphasic behavior, with 60 % of the aa-tRNA$^{Ile}$ hydrolyzed at the rate constants of 2.3 ± 0.3 and 2.4 ± 0.3, respectively. The fast phase, reported in the table, is analogous in value to the previously measured rates for other IleRSs [2, 3]

The values represent the average value ± SEM of three independent experiments.

**Supplementary Table S3. Formation of [$^{32}$P]-AMP and aa-[$^{32}$P]-tRNA$^{Ile}$ using WT and post-transfer editing defective variants [a]**

| IleRS | **Substrate** | $k_{AMP}$ / s$^{-1}$ | $k_{aa\text{-}tRNA}$ / s$^{-1}$ | $k_{AMP}$ / $k_{aa\text{-}tRNA}$ |
|---|---|---|---|---|
| PmIleRS1 WT | L-Ile | 0.578 ± 0.080 | 0.439 ± 0.038 | 1.32 |
| | L-Val | 0.440 ± 0.048 | 0.048 ± 0.006 | 9.17 |
| | L-Nva | 0.429 ± 0.024 | 0.009 ± 0.004 | 47.8 |
| PmIleRS1 D333A/T234R | L-Ile | 0.254 ± 0.008 | 0.186 ± 0.006 | 1.37 |
| | L-Val | 0.244 ± 0.010 | 0.178 ± 0.017 | 1.37 |
| | L-Nva | 0.198 ± 0.015 | 0.121 ± 0.008 | 1.64 |
| PmIleRS2 WT | L-Ile | 0.827 ± 0.061 | 0.982 ± 0.050 | 0.84 |
| | L-Val | 0.534 ± 0.033 | 0.127 ± 0.003 | 4.20 |
| | L-Nva | 0.886 ± 0.074 | 0.024 ± 0.006 | 36.9 |
| PmIleRS2 D324A/T225R | L-Ile | 0.473 ± 0.125 | 0.477 ± 0.013 | 0.99 |
| | L-Val | 0.341 ± 0.011 | 0.316 ± 0.035 | 1.08 |
| | L-Nva | 0.503 ± 0.053 | 0.505 ± 0.032 | 1.00 |

[a] measured using PmIleRSs overexpressed in *E. coli*. To ensure better signal to noise ratio, ATP was present at 200 µM, which is 5-fold lower than in the canonical aminoacylation.

The values represent the average value ± SEM of three independent experiments.

**Supplementary Table S4. List of the plasmids used in this study**

| Plasmid | Description | Reference |
|---|---|---|
| pET28b | *E. coli* protein expression vector, kanamycin resistance | Novagen |
| pET3a | *E. coli* tRNA expression vector, ampicillin resistance | Novagen |
| pUCTV2 | *E. coli* / *P. megaterium* shuttle vector, ampicillin resistance (*E. coli*), tetracycline resistance (*P. megaterium*), temperature sensitive origin of replication (*P. megaterium*) | [4] |
| pMGBm19 | *E. coli* / *P. megaterium* shuttle vector, ampicillin resistance (*E. coli*), chloramphenicol resistance (*P. megaterium*) | [5] |
| pPT7 | *E. coli* / *P. megaterium* shuttle vector, ampicillin resistance (*E. coli*), tetracycline resistance (*P. megaterium*) | [5] |
| pT7_RNAP | *E. coli* / *P. megaterium* shuttle vector, ampicillin resistance (*E. coli*), chloramphenicol resistance (*P. megaterium*), containing gene for T7 RNA polymerase under $P_{xylA}$ promoter | [5] |
| pET28b_PmIleRS1 | pET28b vector containing *ileS1* ORF under T7 promoter with N-terminal $His_6$ tag, used for expression of PmIleRS1 in *E. coli* | This study |
| pET28b_PmIleRS2 | pET28b vector containing *ileS2* ORF under T7 promoter with N-terminal $His_6$ tag, used for expression of PmIleRS2 in *E. coli* | This study |
| pET3a_PmtRNA$^{Ile}$ | pET3a vector containing tRNA$^{Ile}$ gene, major isoacceptor, under T7 promoter, used for expression of tRNA$^{Ile}$ in *E. coli* | This study |
| pUCTV2_ΔileS1 | pUCTV2 vector containing flanking regions of the *ileS1* gene, used for construction of *ΔileS1* strain | This study |
| pUCTV2_ΔileS2 | pUCTV2 vector containing flanking regions of the *ileS2* gene, used for construction of *ΔileS2* strain | This study |
| pP$_{T7}$_PmIleRS1 | pP$_{T7}$ vector containing *ileS1* ORF under T7 promoter with C-terminal $His_6$ tag, used for expression of PmIleRS1 in *P. megaterium* | This study |
| pMGBm19_P$_{xylA}$_PmIleRS2 | pMGBm19 vector containing *ileS2* ORF under xylose-inducible promoter with C-terminal $His_6$ tag, used for expression of PmIleRS2 in *P. megaterium* | This study |
| pMGBm19_P$_{ileS2}$_PmIleRS2 | pMGBm19 vector containing *ileS2* ORF under native promoter and terminator with C-terminal $His_6$ tag | This study |

**Supplementary Table S5. List of the oligonucleotides used in this study**

| **Oligonucleotide name** | **Oligonucleotide sequence (5' – 3')** |
|---|---|
| tRNAIle_S | TCGACTAATACGACTCACTATAGGGCCTATAGCTCAGCTGGTTAGAGCGC ACGCCTGATAAGCGTGAGGTCGGTGGTTCGAGTCCACTTAGGCCCACCAG |
| tRNAIle_A | GATCCTGGTGGGCCTAAGTGGACTCGAACCACCGACCTCACGCTTATCAG GCGTGCGCTCTAACCAGCTGAGCTATAGGCCCTATAGTGAGTCGTATTAG |
| ΔileS1_A | GAGAGAGCGGCCGCAAGACAAAAAGTCCAGCCGTTTATTG |
| ΔileS1_B | GTGTTTTTGTTAATTACTGCTGAACGTAGTGATTTTTTAATG TATCTTTATATTCCATGTTTTACCCTCCAAAAAAAACG |
| ΔileS1_C | TTTTGGAGGGTAAAACATGGAATATAAAGATACATTAAAA AATCACTACGTTCAGCAGTAATTAACAAAAACACGTCCT |
| ΔileS1_D | GAGAGAGGATCCGCTGCACTTGTGAACGTGACCATTC |
| ΔileS2_A | GAGAGAGCGGCCGCAAGACAAAAAGTCCAGCCGTTTATTG |
| ΔileS2_B | AGCTCTGATTTAAAACGCGAAGCCTTACATTCACTTCTTTCATGTGAT ACCACT |
| ΔileS2_C | ACATGAAAGAAGTGAATGTAAGGCTTCGCGTTTTAAATCAGAGCTG |
| ΔileS2_D | GAGAGAGGATCCGTCGTAATGAGGACAGTTGCTG |
| ileS1_F | GAGAGAGGCTAGCATGGAATATAAAGATACATTATTGATGCC |
| ileS1_R | GAGAGAGCTCGAGTTACTGCTGAACGTAGTGATTTTTTAC |
| ileS2_F | GAGAGAGCATATGAAAGAAGTGAATGTAAGGGAGTC |
| ileS2_R | GAGAGAGCTCGAGTCAGCTCTGATTTAAAACGCGAAG |
| ileS1_cp_His_R | GAGAGAGGATCCGGTCTCATTAATTAATGATGATGGTGATG ATGCTGCTGAACGTAGTGATTTTTTAC |
| ileS2_cp_F | GAGAGGGTCTCTCTAGAATGAAAGAAGTGAATGTAAGGGAGTC |
| ileS2_cp_R | GAGAGAGGATCCGGTCTCATTTATCAATGATGATGGTGATGATGGCT CTGATTTAAAACGCGAAG |
| ileS2_cp_pro_F | GAGAGAGCTCGAGTTTCCCTCAGAGCATCCC |
| ileS2_cp_ter_F | GAGAGAGAGCTCGATAAAACCATCACGCTGAAGC |
| ileS2_cp_ter_R | GAGAGAGGTCTCATAAAAAGCACTTACCTCATAGCGG |
| ileS1_T234R_S [a] | AAATTCATCATTTGGACGACAA**G**ACCATGGACAATGCC |
| ileS1_T234R_A [a] | GGCATTGTCCATGGT**C**TTGTCGTCCAAATGATGAATTT' |
| ileS1_D333A_S [a] | GGACACGGGGAAGACG**C**CTTTATCGTTGGTCAA |
| ileS1_D333A_A [a] | TTGACCAACGATAAAG**G**CGTCTTCCCCGTGTCC |
| ileS2_T225R_S [a] | GGGCTGGACGACAA**G**GCCTTGGACG |
| ileS2_T225R_A [a] | CGTCCAAGGC**C**TTGTCGTCCAGCCC |
| ileS2_D324A_S [a] | CCTGCTTACGGGGAAGATG**C**TTACAGAGTTGTAAAAGAA |
| ileS2_D324A_A [a] | TTCTTTTACAACTCTGTAA**G**CATCTTCCCCGTAAGCAGG |

[a] oligonucleotides used for mutagenesis. Bolded red letter indicates substituted nucleotide

### Supplementary Table S6. Construction of the plasmids used in this study

| Plasmid | Construction |
|---|---|
| pET3a_PmtRNA$^{Ile}$ | Oligonucleotides tRNAIle_S/tRNAIle_A were hybridized and cloned in vector pET3a via BamHI/SalI sites. |
| pUCTV2_ΔileS1<br>pUCTV2_ΔileS2 | Flanking regions of *P. megaterium ileS1* gene were amplified from genomic DNA using primers ΔileS1_A/ΔileS1_B (upstream) and ΔileS1_C/ΔileS1_D (downstream). PCR products were fused using SOE-PCR with primers ΔileS1_A/ΔileS1_D and cloned in pUCTV2 via NotI/BamHI sites. Vector pUCTV2_ΔileS2 was created analogously. |
| pET28b_PmIleRS1<br>pET28b_PmIleRS2 | *P. megaterium ileS1* and *ileS2* ORFs were amplified from genomic DNA using primers ileS1_F/ileS1_R and ileS2_F/ileS2_R and cloned in vector pET28b via NheI/XhoI and NdeI/XhoI sites, respectively. |
| $pP_{T7}$_PmIleS1 | *P. megaterium ileS1* ORF was amplified from genomic DNA using primers ileS1_F and ileS1_cp_His_R and cloned in vector $pP_{T7}$ via SpeI(NheI)/BamHI sites. |
| pMGBm19_$P_{xylA}$_PmIleS2 | *P. megaterium ileS2* ORF was amplified from genomic DNA using primers ileS2_cp_F and ileS2_cp_R and cloned in vector pMGBm19 via SpeI(XbaI)/BamHI sites |
| pMGBm19_$P_{ileS2}$_PmIleS2 | *P. megaterium ileS2* ORF together with putative promoter region (approximately 500 bp upstream of ORF) was amplified from genomic DNA using primers ileS2_cp_pro_F and ileS2_cp_R. Putative terminator region (approximately 250 bp downstream of ORF) was amplified using primers ileS2_cp_ter_F and ileS2_cp_ter_R. PCR products were fused using BsaI enzyme and cloned in pMGBm19 via XhoI/SacI sites |

### Supplementary Table S7. Bacteria doubling time

| Phylum | Species | NCBI genomic accession number | IleRS type [a] | Doubling time / min [b] | Ref |
|---|---|---|---|---|---|
| *Tenericutes* | *Acholeplasma laidlawii* | NZ_LS483439.1 | 1 | 57 | [6] |
| *Tenericutes* | *Acholeplasma morum* | NZ_JAFBBG000000000.1 | 1 | 20,4 | [6] |
| *Tenericutes* | *Mycoplasma arginini* | NZ_LR215044.1 | 1 | 197,4 | [6] |
| *Tenericutes* | *Mycoplasma bovis* | NZ_CP068731.1 | 1 | 91,2 | [7] |
| *Tenericutes* | *Mycoplasma fermentans* | NC_014921.1 | 1 | 37,8 | [6] |
| *Tenericutes* | *Mycoplasma gallisepticum* | NC_018406.1 | 1 | 72,6 | [6] |
| *Tenericutes* | *Mycoplasma genitalium* | NC_000908.2 | 1 | 960 | [8] |
| *Tenericutes* | *Mycoplasma pneumoniae* | NZ_LR214945.1 | 1 | 804 | [9] |
| *Tenericutes* | *Mycoplasma pulmonis* | NZ_LR215008 | 1 | 60 | [10] |
| *Tenericutes* | *Spiroplasma apis* | NC_022998 | 1 | 66 | [11] |
| *Tenericutes* | *Spiroplasma citri* | NZ_CP013197 | 1 | 246 | [11] |
| *Tenericutes* | *Spiroplasma floricola* | NZ_CP025057 | 1 | 66 | [11] |
| *Tenericutes* | *Spiroplasma melliferum* | NZ_AMGI01000001 | 1 | 90 | [11] |
| *Tenericutes* | *Spiroplasma mirum* | NZ_CP002082 | 1 | 480 | [11] |
| *Tenericutes* | *Spiroplasma syrphidicola* | NC_021284 | 1 | 60 | [11] |
| *Tenericutes* | *Ureaplasma urealyticum* | NC_011374.1 | 1 | 96 | [12] |
| *Spirochaetes* | *Borrelia burgdorferi* | NZ_CP074054.1 | 2 | 480 | [13] |
| *Spirochaetes* | *Borrelia crocidurae* | NZ_LN609267 | 2 | 420 | [14] |
| *Spirochaetes* | *Borrelia hermsii* | NZ_CP011060.1 | 2 | 270 | [14] |
| *Spirochaetes* | *Borrelia mayonii* | NZ_CP015796.1 | 2 | 540 | [15] |
| *Spirochaetes* | *Borrelia miyamotoi* | NZ_CP021872.1 | 2 | 1020 | [15] |
| *Spirochaetes* | *Borrelia turicatae* | NC_008710.1 | 2 | 420 | [14] |
| *Spirochaetes* | *Brachyspira hyodysenteriae* | NZ_CP046932.1 | 1 | 60 | [16] |
| *Spirochaetes* | *Leptospira interrogans canicola* | NZ_CP020414.2 | 1 | 420 | [17] |
| *Spirochaetes* | *Leptospira interrogans Copenhageni* | NZ_CP020414.2 | 1 | 360 | [18] |
| *Spirochaetes* | *Leptospira interrogans icterohaemorrhagiae* | NZ_CP020414.2 | 1 | 348 | [19] |
| *Spirochaetes* | *Leptospira kobayashii* | NZ_BFBA01000006 | 1 | 480 | [20] |
| *Spirochaetes* | *Leptospira pomona* | NZ_CP020414.2 | 1 | 840 | [17] |
| *Spirochaetes* | *Treponema denticola* | NC_002967.9 | 2 | 840 | [21] |
| *Spirochaetes* | *Treponema pallidum* | NC_016842.1 | 2 | 1800 | [22] |

| | | | | | |
|---|---|---|---|---|---|
| *Deinococcus-Thermus* | *Deinococcus deserti* | NC_012526 | 2 | 180 | [23] |
| *Deinococcus-Thermus* | *Deinococcus radiodurans* | NZ_CP031500.1 | 2 | 130 | [24] |
| *Deinococcus-Thermus* | *Deinococcus sp AJ005* | CP044990.1 | 2 | 231 | [25] |
| *Deinococcus-Thermus* | *Thermus thermophilus* | NC_006461.1 | 2 | 150 | [26] |
| *Cyanobacteria* | *Microcystis aeruginosa* | NZ_JXYX01000003 | 1 | 2160 | [27] |
| *Cyanobacteria* | *Synechococcus elongatus* | NZ_CP033061 | 1 | 114 | [28] |
| *Cyanobacteria* | *Crocosphaera watsonii* | NZ_AADV02000003.1 | 1 | 1980 | [29] |
| *Cyanobacteria* | *Gloeothece sp. ATCC 27152* | CP001291 | 1 | 4140 | [30] |
| *Cyanobacteria* | *Cylindrospermopsis raciborskii* | NZ_CP073250 | 1 | 2935,8 | [31] |
| *Cyanobacteria* | *Nostoc commune* | NZ_BDUD01000001 | 1 | 18288 | [32] |
| *Cyanobacteria* | *Aphanizomenon flos-aquae* | CP051188 | 1 | 3696,6 | [31] |
| *Cyanobacteria* | *Prochlorococcus marinus* | NC_005042 | 1 | 1440 | [33] |
| *Cyanobacteria* | *Acaryochloris marina* | NC_009925 | 1 | 3300 | [34] |
| *Cyanobacteria* | *Aphanizomenon aphanizomenoides* | NZ_LUFH02000003 | 1 | 2772,6 | [31] |
| *Cyanobacteria* | *Gloeobacter violaceus* | NC_005125.1 | 1 | 4380 | [35] |
| *Cyanobacteria* | *Synechocystis sp PCC 6803* | CP073017 | 1 | 307,8 | [36] |
| *Cyanobacteria* | *Trichodesmium erythraeum* | JAGGDU010000202 | 1 | 2495,4 | [37] |
| *Chlamydiae* | *Chlamydia trachomatis* | NC_000117 | 2 | 108 | [38] |
| *Chlamydiae* | *Chlamydia pneumonia* | NC_005043 | 2 | 204 | [39] |
| *Bacteroidetes* | *Porphyromonas gingivalis* | NC_010729.1 | 2 | 324 | [40] |
| *Bacteroidetes* | *Prevotella intermedia* | NZ_CP024728 | 2 | 142,2 | [41] |
| *Bacteroidetes* | *Tannerella forsythia* | NZ_FMML01000083 | 2 | 122,4 | [42] |
| *Bacteroidetes* | *Bacteroides caccae* | NZ_VVYG01000008 | 2 | 47,1 | [43] |
| *Bacteroidetes* | *Bacteroides fragilis* | NZ_CP069563.1 | 2 | 60,4 | [43] |
| *Bacteroidetes* | *Bacteroides ovatus* | NZ_CP012938 | 2 | 78,8 | [43] |
| *Bacteroidetes* | *Bacteroides thetaiotaomicron* | NZ_CP040530.1 | 2 | 46 | [43] |
| *Bacteroidetes* | *Bacteroides uniformis* | NZ_CP072255 | 2 | 47,9 | [43] |
| *Bacteroidetes* | *Parabacteroides distasonis* | NZ_CP050956 | 2 | 55,2 | [43] |
| *Actinobacteria* | *Actinomyces naeslundii* | NZ_CP066049 | 2 | 130,2 | [42] |
| *Actinobacteria* | *Bifidobacterium animalis* | NC_017216 | 2 | 158 | [44] |
| *Actinobacteria* | *Bifidobacterium bifidum* | NZ_AKCA01000001 | 2 | 88 | [45] |
| *Actinobacteria* | *Bifidobacterium breve* | NZ_CP006712 | 2 | 118 | [45] |
| *Actinobacteria* | *Bifidobacterium longum* | NC_015052 | 2 | 192 | [44] |

| | | | | | |
|---|---|---|---|---|---|
| *Actinobacteria* | *Bifidobacterium pseudolongum* | NZ_CP022544 | 2 | 174 | [44] |
| *Actinobacteria* | *Corynebacterium diphtheriae* | NZ_LN831026 | 2 | 70 | [46] |
| *Actinobacteria* | *Corynebacterium glutamicum* | NZ_LOQW01000006 | 2 | 131,4 | [47] |
| *Actinobacteria* | *Cutibacterium acnes* | NC_021085 | 2 | 65,3 | [43] |
| *Actinobacteria* | *Micrococcus luteus* | NZ_CP082331 | 2 | 77,4 | [43] |
| *Actinobacteria* | *Mycobacterium avium* | NZ_AUZQ01000023 | 2 | 204 | [48] |
| *Actinobacteria* | *Mycobacterium chelonea* | NZ_CP007220 | 2 | 300 | [49] |
| *Actinobacteria* | *Mycobacterium fortuitum* | NZ_CP011269 | 2 | 240 | [49] |
| *Actinobacteria* | *Mycobacterium leprae* | NZ_CP029543 | 2 | 14400 | [50] |
| *Actinobacteria* | *Mycobacterium marinum* | NZ_CP058277 | 2 | 534 | [51] |
| *Actinobacteria* | *Mycobacterium smegmatis* | NZ_CP054795 | 2 | 120 | [52] |
| *Actinobacteria* | *Mycobacterium tuberculosis* | NC_000962 | 2 | 882 | [53] |
| *Actinobacteria* | *Streptomyces aureofaciens* | NZ_JPRF03000032 | 2 | 231 | [54] |
| *Actinobacteria* | *Streptomyces coelicolor* | NC_003888 | 2 | 132 | [55] |
| *Actinobacteria* | *Streptomyces lividans* | CP071800 | 2 | 109,2 | [56] |
| *Actinobacteria* | *Streptomyces thermoviolaceus* | NZ_BMVZ01000002 | 2 | 237,6 | [57] |
| *Actinobacteria* | *Tropheryma whipplei* | NC_004551 | 2 | 2040 | [58] |
| *Proteobacteria* | *Agrobacterium tumefaciens* | NZ_CP033031 | 1 | 150 | [59] |
| *Proteobacteria* | *Anaplasma phagocytophilum* | NC_021880 | 2 | 1464 | [60] |
| *Proteobacteria* | *Bradyrhizobium japonicum* | NZ_CP058354 | 1 | 390 | [61] |
| *Proteobacteria* | *Brucella melitensis* | NC_003318 | 1 | 150 | [62] |
| *Proteobacteria* | *Brucella suis* | NC_004311 | 1 | 360 | [63] |
| *Proteobacteria* | *Caulobacter crescentus* | NC_011916 | 1 | 89 | [64] |
| *Proteobacteria* | *Ehrlichia canis* | NC_007354 | 2 | 1560 | [60] |
| *Proteobacteria* | *Ehrlichia chaffeensis* | NZ_CP007480 | 2 | 1410 | [60] |
| *Proteobacteria* | *Mesorhizobium loti* | NZ_QGGH01000002 | 1 | 240 | [65] |
| *Proteobacteria* | *Rhodopseudomonas palustris* | NZ_CP066699 | 1 | 504 | [66] |
| *Proteobacteria* | *Rickettsia bellii* | NZ_CP015010 | 2 | 480 | [67] |
| *Proteobacteria* | *Rickettsia conorii* | NC_003103 | 2 | 540 | [68] |
| *Proteobacteria* | *Rickettsia helvetica* | NZ_CM001467 | 2 | 1200 | [69] |
| *Proteobacteria* | *Rickettsia prowazekii* | NC_017049 | 2 | 480 | [70] |
| *Proteobacteria* | *Rickettsia rickettsii* | NC_010263 | 2 | 540 | [68] |
| *Proteobacteria* | *Rickettsia slovaca* | NC_016639 | 2 | 1218 | [69] |
| *Proteobacteria* | *Rickettsia typhi* | NC_017066 | 2 | 936 | [69] |
| *Proteobacteria* | *Sinorhizobium meliloti* | NC_020528 | 1 | 140 | [71] |
| *Proteobacteria* | *Wolbachia pipientis* | NZ_CP050531 | 2 | 840 | [72] |

| | | | | | |
|---|---|---|---|---|---|
| *Proteobacteria* | *Achromobacter denitrificans* | NZ_CP053986 | 1 | 43,1 | [43] |
| *Proteobacteria* | *Bordetella bronchiseptica* | NZ_LR134326 | 1 | 108 | [73] |
| *Proteobacteria* | *Bordetella hinzii* | NZ_CP021395 | 1 | 114 | [73] |
| *Proteobacteria* | *Bordetella parapertussis* | NC_018828 | 1 | 168 | [74] |
| *Proteobacteria* | *Bordetella petrii* | NC_010170 | 1 | 114 | [73] |
| *Proteobacteria* | *Chromobacterium violaceum* | NZ_CP069587 | 1 | 74,76 | [75] |
| *Proteobacteria* | *Neisseria gonorrhoeae* | NZ_AP023069 | 1 | 60 | [76] |
| *Proteobacteria* | *Neisseria lactamica* | NZ_CP031253 | 1 | 80 | [77] |
| *Proteobacteria* | *Neisseria meningitidis* | NZ_CP021520 | 1 | 40 | [77] |
| *Proteobacteria* | *Nitrosomonas europaea* | NC_004757 | 1 | 405 | [78] |
| *Proteobacteria* | *Ralstonia solanacearum* | NZ_CP012943 | 1 | 111,6 | [79] |
| *Proteobacteria* | *Geobacter sulfurreducens* | NC_002939 | 1 | 415,8 | [80] |
| *Proteobacteria* | *Campylobacter jejuni* | NC_002163 | 1 | 90 | [81] |
| *Proteobacteria* | *Helicobacter hepaticus* | NC_004917 | 1 | 252 | [82] |
| *Proteobacteria* | *Helicobacter pylori* | NZ_CP071982 | 1 | 50 | [83] |
| *Proteobacteria* | *Wolinella succinogenes* | NC_005090 | 1 | 60 | [84] |
| *Proteobacteria* | *Acinetobacter baumannii* | NZ_CP043953 | 1 | 48 | [85] |
| *Proteobacteria* | *Citrobacter rodentium* | NC_013716 | 1 | 66 | [86] |
| *Proteobacteria* | *Coxiella burnetii* | NC_002971 | 1 | 546 | [87] |
| *Proteobacteria* | *Enterobacter cloacae* | NZ_CP009756 | 1 | 74,8 | [43] |
| *Proteobacteria* | *Escherichia coli* | NC_000913 | 1 | 25 | [88] |
| *Proteobacteria* | *Haemophilus ducreyi* | NZ_CP015425 | 1 | 120 | [89] |
| *Proteobacteria* | *Haemophilus influenzae* | NZ_QWLX01000001 | 1 | 30 | [90] |
| *Proteobacteria* | *Klebsiella pneumoniae* | NC_016845 | 1 | 38 | [91] |
| *Proteobacteria* | *Pasteurella multocida* | NZ_CP028926 | 1 | 30 | [92] |
| *Proteobacteria* | *Photorhabdus luminescens* | NZ_JXSK01000004 | 1 | 126 | [93] |
| *Proteobacteria* | *Proteus hauseri* | NZ_CP026364 | 1 | 104,6 | [43] |
| *Proteobacteria* | *Proteus mirabilis* | NC_010554 | 1 | 80,7 | [43] |
| *Proteobacteria* | *Pseudomonas aeruginosa* | NC_002516 | 1 | 30 | [94] |
| *Proteobacteria* | *Pseudomonas fluorescens* | NZ_LT907842 | 1 | 33 | [95] |
| *Proteobacteria* | *Pseudomonas putida* | NC_021505 | 1 | 54,6 | [96] |
| *Proteobacteria* | *Pseudomonas syringae* | NZ_CP068034 | 1 | 76,2 | [97] |
| *Proteobacteria* | *Salmonella enterica* | NC_003197 | 1 | 32 | [98] |
| *Proteobacteria* | *Shewanella oneidensis* | NC_004347 | 1 | 40 | [99] |
| *Proteobacteria* | *Vibrio cholerae* | NZ_AP014524 | 1 | 16 | [100] |
| *Proteobacteria* | *Vibrio parahaemolyticus* | NC_004603 | 1 | 12 | [100] |

| | | | | | |
|---|---|---|---|---|---|
| *Proteobacteria* | *Vibrio vulnificus* | NZ_CP016321 | 1 | 18 | [100] |
| *Proteobacteria* | *Xanthomonas arboricola* | NZ_HG999362 | 1 | 91,8 | [97] |
| *Proteobacteria* | *Xylella fastidiosa* | NC_004556 | 1 | 307,8 | [101] |
| *Proteobacteria* | *Yersinia pestis* | NC_017168 | 1 | 75 | [102] |
| *Proteobacteria* | *Yersinia pseudotuberculosis* | NZ_CP009712 | 1 | 152,6 | [43] |
| *Firmicutes* | *Bacillus amyloliquefaciens* | NC_020272 | 1 | 22 | [88] |
| *Firmicutes* | *Bacillus anthracis* | NC_007530 | 1 & 2 | 30 | [88] |
| *Firmicutes* | *Bacillus cereus* | NZ_CP072774 | 1 & 2 | 18 | [103] |
| *Firmicutes* | *Bacillus circulans* | NZ_CP053989 | 1 & 2 | 38 | [88] |
| *Firmicutes* | *Bacillus clausii* | NZ_NPBN01000012 | 1 | 37 | [88] |
| *Firmicutes* | *Bacillus cohnii* | NZ_CP018866 | 1 | 29 | [88] |
| *Firmicutes* | *Bacillus flexus* | NZ_JAEMWV010000001 | 1 | 20 | [88] |
| *Firmicutes* | *Bacillus glycinifermentans* | NZ_CP035232 | 1 | 28 | [88] |
| *Firmicutes* | *Bacillus lentus* | NZ_LS483476 | 1 | 26 | [88] |
| *Firmicutes* | *Bacillus licheniformis* | NZ_CP014842 | 1 | 21 | [88] |
| *Firmicutes* | *Priestia megaterium* | CP009920 | 1 & 2 | 22 | [88] |
| *Firmicutes* | *Bacillus pumilus* | NZ_PVQT01000001 | 1 | 20 | [88] |
| *Firmicutes* | *Bacillus shackletonii* | NZ_LJJC01000004 | 1 & 2 | 30 | [88] |
| *Firmicutes* | *Bacillus subtilis* | NC_000964 | 1 | 22 | [88] |
| *Firmicutes* | *Bacillus thuringiensis* | NZ_CM000753 | 1 & 2 | 20 | [103] |
| *Firmicutes* | *Brevibacillus laterosporus* | NZ_CP017705 | 1 & 2 | 28 | [88] |
| *Firmicutes* | *Carnobacterium maltaromaticum* | NZ_JAGYWR010000002 | 1 | 68,6 | [43] |
| *Firmicutes* | *Enterococcus caccae* | NZ_KB946333 | 1 | 60 | [88] |
| *Firmicutes* | *Enterococcus durans* | NZ_JAAMRZ010000006 | 1 | 38 | [88] |
| *Firmicutes* | *Enterococcus faecalis* | NZ_KB944666 | 1 | 48 | [104] |
| *Firmicutes* | *Enterococcus faecium* | NZ_CP038996 | 1 | 30 | [88] |
| *Firmicutes* | *Enterococcus mundtii* | NZ_CP018061 | 1 | 30 | [88] |
| *Firmicutes* | *Lactobacillus acidophilus* | NC_021181 | 1 | 74 | [43] |
| *Firmicutes* | *Lactobacillus fermentum* | NZ_BJLV01000021 | 1 | 91,3 | [43] |
| *Firmicutes* | *Lactobacillus gasseri* | NZ_CP072178 | 1 | 95,3 | [43] |
| *Firmicutes* | *Lactobacillus paracasei* | NC_014334 | 1 | 88,5 | [43] |
| *Firmicutes* | *Lactobacillus plantarum* | NZ_CP030105 | 1 | 32 | [88] |
| *Firmicutes* | *Lactobacillus reuteri* | NZ_CP045049 | 1 | 36,6 | [105] |
| *Firmicutes* | *Lactobacillus rhamnosus* | NZ_PKJX01000001 | 1 | 76,1 | [43] |
| *Firmicutes* | *Lactobacillus ruminis* | NZ_GL833109 | 1 | 179,1 | [43] |
| *Firmicutes* | *Lactococcus lactis* | NC_020450 | 1 | 30 | [88] |

| | | | | | |
|---|---|---|---|---|---|
| *Firmicutes* | *Listeria innocua* | NZ_JACTKR010000004 | 1 | 60 | [106] |
| *Firmicutes* | *Listeria monocytogenes* | NC_003210 | 1 | 38 | [88] |
| *Firmicutes* | *Lysinibacillus sphaericus* | NZ_CP019980 | 1 | 23 | [88] |
| *Firmicutes* | *Oceanobacillus oncorhynchi* | NZ_CDGG01000001 | 1 | 70 | [88] |
| *Firmicutes* | *Oceanobacillus sojae* | NZ_BJYM01000002 | 1 & 2 | 50 | [88] |
| *Firmicutes* | *Paenibacillus graminis* | NZ_CP009287 | 2 | 94 | [88] |
| *Firmicutes* | *Paenibacillus macerans* | NZ_UGSI01000001 | 2 | 25 | [88] |
| *Firmicutes* | *Paenibacillus polymyxa* | NZ_CP040829 | 2 | 70 | [88] |
| *Firmicutes* | *Pediococcus pentosaceus* | NC_008525 | 1 | 35 | [88] |
| *Firmicutes* | *Rummeliibacillus pycnus* | NZ_KZ614145 | 1 | 38 | [88] |
| *Firmicutes* | *Rummeliibacillus stabekisii* | NZ_BJVD01000001 | 1 | 30 | [88] |
| *Firmicutes* | *Sporosarcina luteola* | NZ_BJYL01000004 | 1 | 65 | [88] |
| *Firmicutes* | *Staphylococcus aureus* | NC_007795 | 1 | 24 | [107] |
| *Firmicutes* | *Staphylococcus capitis* | NZ_CP007601 | 1 | 50 | [88] |
| *Firmicutes* | *Staphylococcus epidermidis* | NZ_CP035288 | 1 | 17 | [108] |
| *Firmicutes* | *Streptococcus agalactiae* | NZ_CP012480 | 1 | 35 | [109] |
| *Firmicutes* | *Streptococcus infantarius* | NZ_JAHCZQ010000002 | 1 | 30 | [88] |
| *Firmicutes* | *Streptococcus mutans* | NZ_CP044221 | 1 | 66 | [110] |
| *Firmicutes* | *Streptococcus pneumoniae* | NZ_CP020549 | 1 | 30 | [111] |
| *Firmicutes* | *Streptococcus pyogenes* | NZ_CP010450 | 1 | 40 | [112] |
| *Firmicutes* | *Blautia producta* | NZ_CP039126 | 2 | 47,3 | [43] |
| *Firmicutes* | *Caldanaerobacter subterraneus* | NC_003869 | 1 | 65 | [113] |
| *Firmicutes* | *Clostridium acetobutylicum* | NC_015687 | 2 | 90 | [114] |
| *Firmicutes* | *Clostridium bifermentans* | NZ_CP079737 | 2 | 14,8 | [115] |
| *Firmicutes* | *Clostridium botulinum* | NC_009495 | 2 | 60 | [116] |
| *Firmicutes* | *Clostridium cadaveris* | NZ_OBJM01000005 | 2 | 31,3 | [115] |
| *Firmicutes* | *Clostridium difficile* | NZ_CP076401 | 2 | 40 | [117] |
| *Firmicutes* | *Clostridium indolis* | NZ_AZUI01000001 | 2 | 142,5 | [43] |
| *Firmicutes* | *Clostridium innocuum* | NZ_CP048838 | 2 | 44,1 | [115] |
| *Firmicutes* | *Clostridium kluyveri* | NC_011837 | 2 | 173,4 | [118] |
| *Firmicutes* | *Clostridium perfringens* | NC_008261 | 2 | 12 | [119] |
| *Firmicutes* | *Clostridium tetani* | NZ_QMAX01000007 | 2 | 90,2 | [120] |
| *Firmicutes* | *Clostridium thermobutyricum* | NZ_KB850956 | 2 | 32 | [121] |
| *Firmicutes* | *Enterocloster clostridioformis* | NZ_KB850969 | 2 | 84,1 | [43] |
| *Firmicutes* | *Erysipelatoclostridium ramosum* | NZ_CP068170 | 1 | 32,2 | [115] |

[a] IleRSs were classified as type 1 or type 2 based of the phylogenetic analysis

[b] doubling time is the minimal published doubling time for the best growth conditions

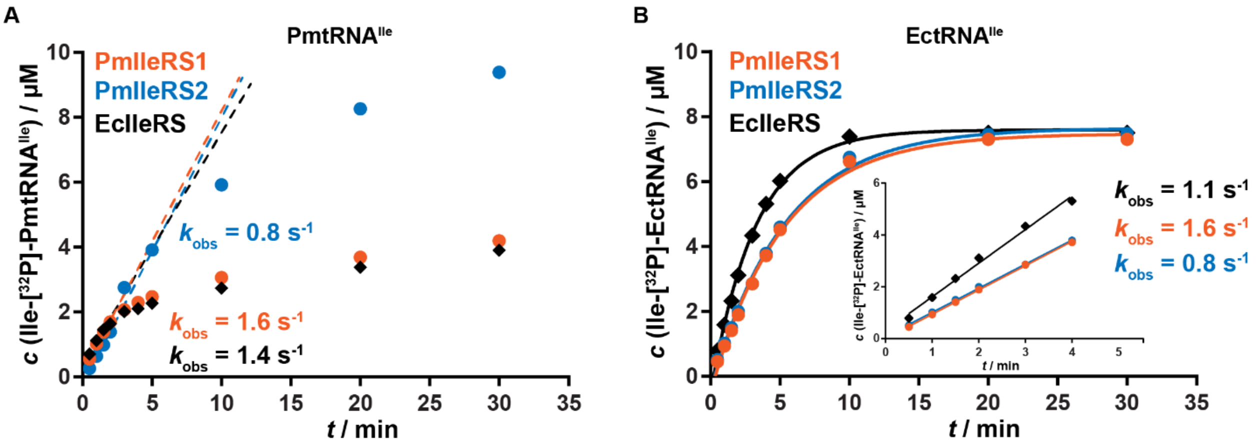


**Supplementary Figure S1. Biphasic nature of PmtRNA$^{Ile}$ aminoacylation profiles**

A) 15 µM active PmtRNA$^{Ile}$ overexpressed in *E. coli* was aminoacylated using PmIleRS1 (10 nM) or PmIleRS2 (20 nM) and also EcIleRS as a control (♦, 20 nM). The time courses were biphasic, which was less pronounced for PmIleRS2. PmIleRS1 and PmIleRS2, as well as EcIleRS, aminoacylated about 10 % of PmtRNA$^{Ile}$ with the rates ($k_{obs}$) that match the aminoacylation rates of these enzymes with EctRNA$^{Ile}$ (panel B, note lack of a biphasic character). The kinetically productive fast aminoacylation phase (indicated with dotted lines) was used for obtaining the steady-state parameters (**Table 2** and **Supplementary Table S3**). The origin of biphasic profiles only with PmtRNA$^{Ile}$ is still not clear.

B) Aminoacylation of 10 µM active EctRNA$^{Ile}$ overexpressed in *E. coli* did not display a biphasic behavior with neither of IleRSs. EcIleRS and [$^{32}$P]-EctRNA$^{Ile}$ were produced as previously published [122].

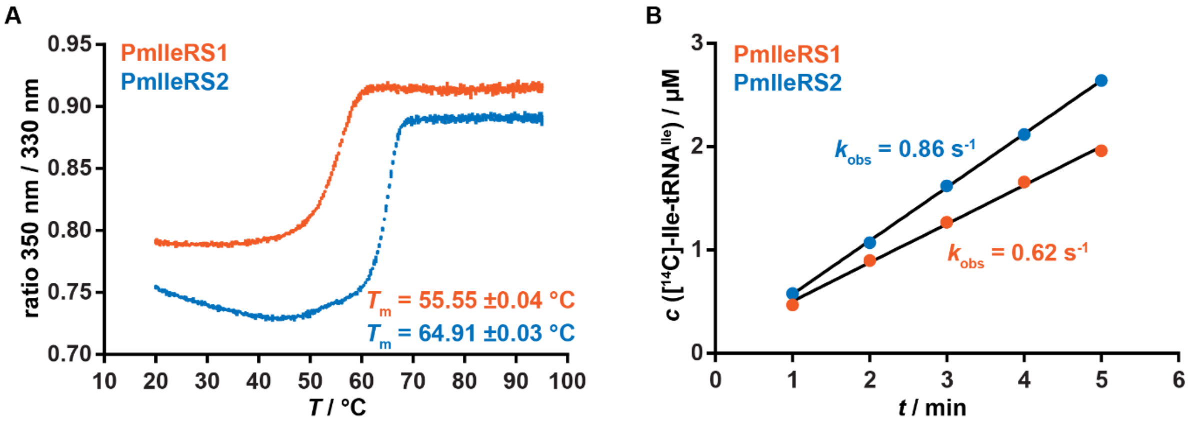


**Supplementary Figure S2. PmIleRSs stability and aminoacylation activity after prolonged preincubation**

A) Thermal unfolding of PmIleRSs measured using Nanotemper Prometheus NT.48. Apparent thermal melting temperature ($T_m$) was calculated from the ratio of tryptophan emission at 350 nm and 330 nm. The values represent the average value ± standard deviation of three independent experiments and error bars represent standard deviation.

B) The enzymes were preincubated with $tRNA^{Ile}$ and ATP for 2.5 h at 37 °C prior aminoacylation at 30 °C. Reactions were started by adding isoleucine (final 50 µM $[^{14}C]$-isoleucine). The observed aminoacylation rate constant ($k_{obs}$) was determined from the slope of the product formation in time divided by the enzyme concentration. No significant loss of enzyme's activity due to prolonged incubation was observed.

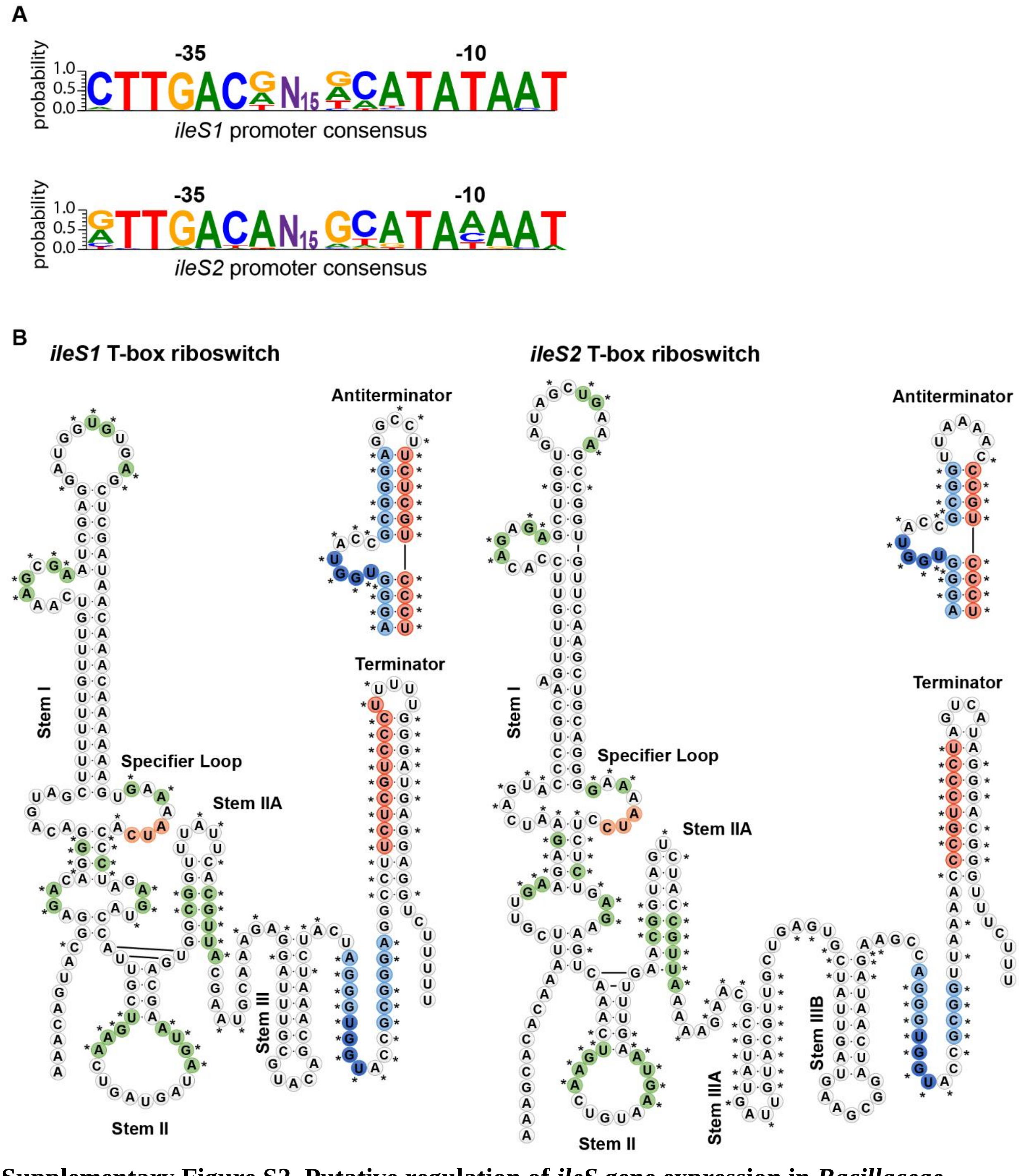


**Supplementary Figure S3. Putative regulation of *ileS* gene expression in *Bacillaceae***

A) Bioinformatic analysis of the upstream regions of 90 *ileS1* and 32 *ileS2* genes from *Bacillaceae* generated the consensus promoter sequences. The consensus *iles1* promoter entails the conserved canonical promoter sequences, suggesting its recognition by canonical $\sigma^{70}$ subunits of bacterial

RNA polymerase. In sharp contrast, the promoter of *ileS2* lacks the relevant conserved elements of the canonical promoters indicating its inducible character. N15 depicts average distance of 15 nucleotides between -35 and -10 promoter regions.

B) A secondary structure model of *P. megaterium ileS* T-Box riboswitch leader region. Structural elements conserved among known T-Box riboswitches are labeled (stem I, II, IIA, III). PmIleRS2 riboswitch stem III is split into two stems (stem IIIA and stem IIIB). Orange nucleotides in the specifier sequence correspond to the isoleucine codon and interact with the tRNA$^{Ile}$ anticodon. The red and blue nucleotides can interact to form antiterminator conformation which exposes UGGU nucleotides (dark blue) for interaction with the uncharged 3’-end of tRNA$^{Ile}$, enabling transcriptional read-through. 90 *ileS1* and 32 *ileS2* leader regions upstream of *ileS* genes were aligned and the conserved (> 90 %) nucleotides in all analyzed sequences are marked as green. Nucleotides conserved only in *ileS1* or *ileS2* leader regions are marked with asterisk (*).

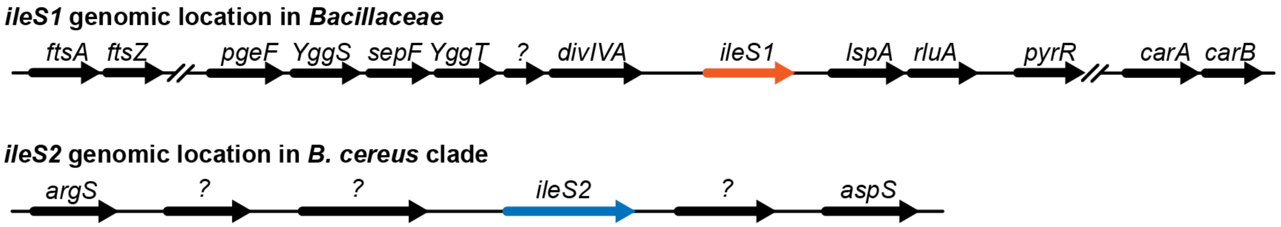


**Supplementary Figure S4. Genomic location of *ileS* genes in *Bacillaceae***

*ileS1* gene is always present at the same, well-established location within the genome of *Bacillaceae*. A specific location for *ileS2* gene was not found, suggesting multiple unrelated gene transfers. The exception is the Cereus clade where *ileS2* seems to reside at a defined genome location.

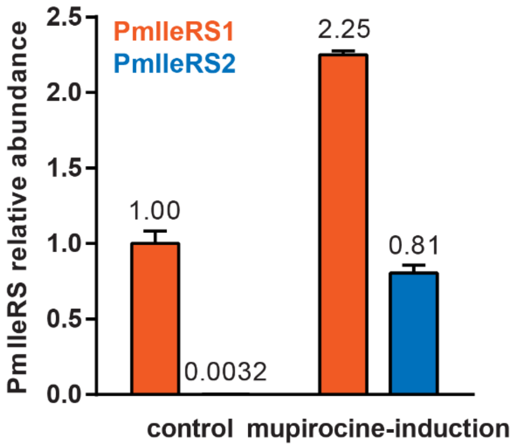


**Supplementary Figure S5. Relative abundance of PmIleRSs.**

The main drawback of *P. megaterium* whole-proteome analysis (**Figure 5**) concerns close to the detection limit expression of PmIleRS2 under the control (mupirocin-free) conditions. To reduce the sample complexity, and improve detection of PmIleRS2, the same proteome samples used for the whole-proteome analysis (control plus mupirocin-induction) were resolved on SDS-PAGE and the bands corresponding to molecular weight of approximately 90 000 – 130 000 were cut and subjected to in-gel digestion (for details see Supplementary materials and methods) prior to MS analysis. Inspection of the processed data revealed that the sequence coverage of PmIleRS2 under the control conditions improved significantly compared to whole proteome MS analysis. Because it was not possible to use LFQ or conventional, whole sample normalization, 41 proteins of high abundance and confidence were selected for sample normalization. Relative abundance is set to 1.0 for PmIleRS1 grown under the control (mupirocin-free) conditions. Error bars represent standard deviation. PmIleRS2 abundance increased up to 250-fold upon mupirocin treatment, lagging slightly behind the PmIleRS1 abundance under the control conditions.

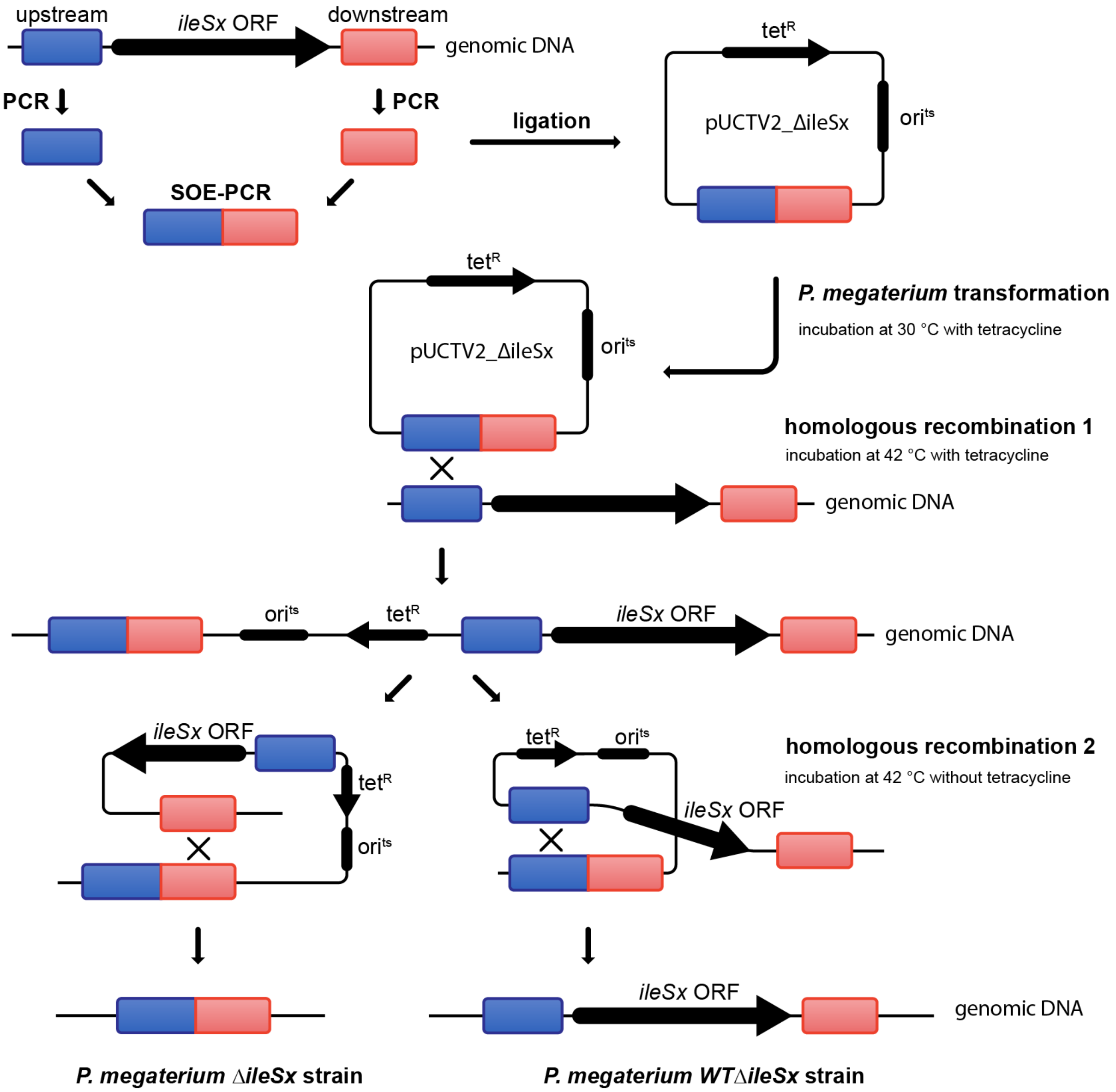


**Supplementary Figure S6. Schematic presentation of the *P. megaterium ΔileSx* strain design.** Flanking regions approximately 1 kb upstream (blue) and downstream (red) of *ileSx* ORF were amplified from genomic DNA, fused using overlap extension PCR (OE-PCR) and cloned in pUCTV2 shuttle vector containing tetracycline resistance ($tet^R$) for selection in *P. megaterium* and temperature sensitive origin of replication ($ori^{ts}$). First and last 7 amino acids of *ileSx* ORF were preserved and fused in frame to eliminate polar effects on adjacent genes. After selecting freely replicating plasmid on tetracycline plates at permissive temperature (30 °C), temperature was raised to 42 °C (nonpermissive temperature) to select genomic integration of the plasmid *via* homologous recombination mediated by flanking regions. At nonpermissive temperature, replication of the pUCTV2 plasmid is arrested thus only viable colonies are the ones with pUCTV2 integrated in genome. Second recombination event was screened by incubating several colonies at 42 °C without tetracycline thus enabling excision of the plasmid from the genome. This event can lead to a knockout genotype (designated as *ΔileSx* strain) if the second recombination event happened at the place opposite of the first recombination event (e.i. if the first recombination happened *via* upstream flanking region, then second needs to happen *via* downstream flanking region) or a wild-type genotype (designated as *WTΔileSx* strain) if the second recombination event happened at the same place as the first one. Further cultivation of strains at nonpermissive temperature without tetracycline ultimately resulted in plasmid curing and loss of tetracycline resistance. Curing of plasmid pUCTV2_ΔileS1 was facilitated by addition of 1 μM mupirocin. At least two biological replicates (obtained *via* two separate recombination events) were obtained for each strain, and all strains were sequenced using primers outside flanking regions to confirm no mutations or indels occurred. X denotes number corresponding to *ileS1* or *ileS2*.

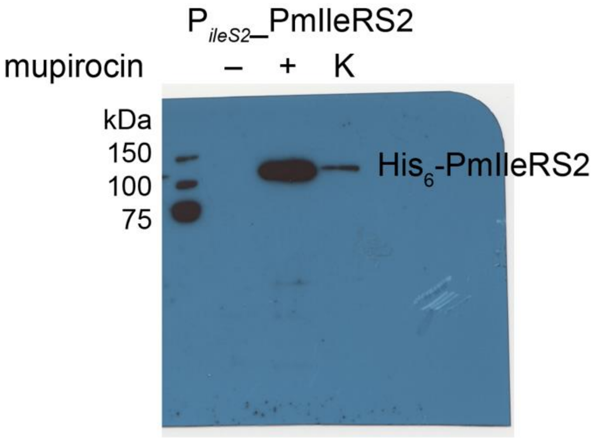


**Supplementary Figure S7. Mupirocin-induced PmIleRS2 expression in *ΔileS2* strain.**

Plasmid pMGBm19_P$_{ileS2}$_PmIleRS2 (having PmIleRS2 under its putative native promoter) was transformed into *ΔileS2* strain. The cultures were grown in LB medium, supplemented with chloramphenicol, until mid-exponential phase in the presence (+) and absence (–) of 10 µM mupirocin. Twenty micrograms of total cellular proteins were applied per line. Purified His$_6$-PmIleRS2 was used as a control (K) and signals were detected using anti-His$_6$ antibodies.

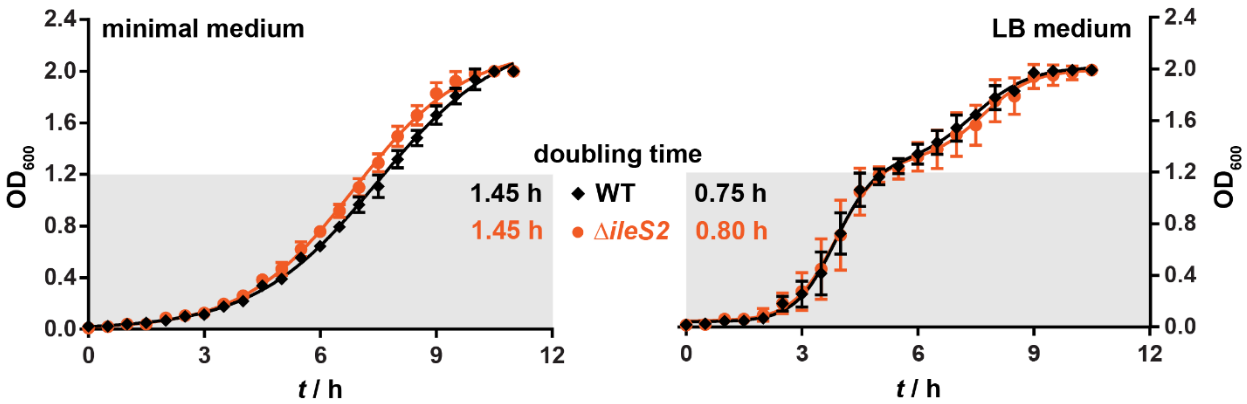


**Supplementary Figure S8. Growth analysis of *P. megaterium* strains.**

The growth curves were followed in minimal medium (left, A5 medium with 0.5 % w/v glucose instead of yeast extract [123]) and LB (right). The full profiles were fitted to logistic (left) [124] and biphasic equations (right) (GraphPad Prism). The exponential phases ($OD_{600}$ < 1.2, marked with gray) were separately fitted to $y = y_0 \times e^{r \times t}$, the growth rates ($r$) extrapolated, and the calculated doubling time is depicted. Experiments were performed at least three times using at least two biological replicates. Error bars represent standard deviation.

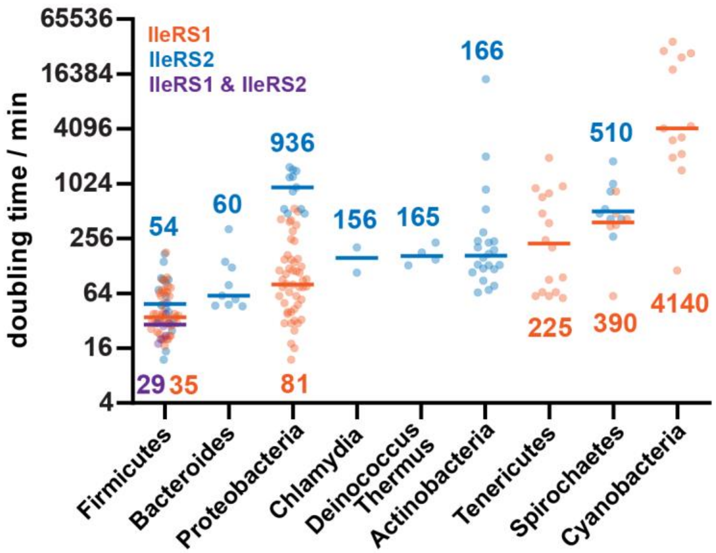


**Supplementary Figure S9. Relationship between bacterial phyla and the doubling time.** Each dot represents one bacterial species grouped by phylum and colored according to the genomic IleRS type (n (IleRS1) = 127, n (IleRS2) = 72, n (IleRS1 & IleRS2) = 8). Line represents median doubling time. All references and accession numbers are provided in **Supplementary Table S7**.

### Supplementary Materials and methods

*Amino acid activation*

Amino acid activation, the first step of two-step overall aminoacylation, was measured using ATP-$PP_i$ exchange assay at 30 °C in a buffer containing 50 mM Hepes-KOH, pH 7.5, 20-30 mM $MgCl_2$, 5 mM DTT, 1 mM [$^{32}$P]-$PP_i$, 0.1 mg/mL BSA and ATP, amino acid and the enzyme. Enzymes were present at 20-100 nM. Steady state kinetic parameters were determined by varying the corresponding substrate from 0.1 – 10 × $K_M$, while the other substrate was kept at the saturating concentration (10 × $K_M$). Effect of tRNA on kinetic parameters was determined by adding 10 µM renatured oxidized $tRNA^{Ile}$ ($tRNA^{Ile}_{ox}$). Reactions were started by addition of the enzyme and stopped by mixing 1.5 µL reaction mixture with 3 µL quench solution (750 mM NaOAc, pH 4.5, 0.15 % SDS). Separation of formed [$^{32}$P]-ATP from [$^{32}$P]-$PP_i$ was performed by thin-layer chromatography on polyethylenimine plates (Macherey-Nagel, Düren, Germany) in a buffer containing 750 mM $KH_2PO_4$, pH 3.5 and 4 M urea. Signal visualization was performed on a Typhoon Phosphorimager (GE Healthcare) and quantified using ImageQuant as described [125]. Kinetic parameters ($K_M$ and $k_{cat}$) were determined by fitting the data directly to Michaelis-Menten equation using GraphPad Prism.

*Mupirocin inhibition of PmIleRS activity*

The mechanism of mupirocin action was studied by measuring inhibition of PmIleRS enzymes in the activation reaction using ATP-$PP_i$ exchange assay. We observed time-dependent inhibition (**Figure 3A**) in the case of PmIleRS1 as well as high mupirocin affinity, both indicating slow, tight-binding inhibition (described in depth in [126, 127]) as previously shown for IleRS1 from *S. aureus*. Tight binding indicates that the concentration of inhibitor used to inhibit the enzyme's activity is in the same concentration range as the free enzyme, thus binding of the inhibitor to the enzyme causes a significant depletion of the free inhibitor concentration. To overcome tight-binding limits, both substrates were present at 60 × $K_M$ (24 mM ATP and 120 µM isoleucine), the enzyme concentration was reduced to 5 nM, and reactions were started by addition of the enzyme. Slow-binding inhibition was indicated by time-dependent inhibition of the PmIleRS1 activity (**Figure 3A**). Thus, time-course data were fitted to equation 1 describing slow-binding inhibition

$$Y = V_s \times t + \frac{(V_0 - V_s) \times (1 - e^{-k \times t})}{k} \text{ (eq. 1)}$$

where $Y$ is the concentration of the formed product ([$^{32}$P]-ATP) in time $t$, $V_0$ is initial velocity, $V_s$ is steady-state velocity, and $k$ is the apparent first-order rate constant for the establishment of the equilibrium between enzyme, inhibitor and enzyme:inhibitor complex. The two most common slow-binding mechanisms involve either slow, one-step formation of the enzyme:inhibitor complex (mechanism I) or initially rapid binding of the inhibitor followed by a slow conformational change of EI to E*I (mechanism II) [126]. Dependence of $k$ vs [I] may distinguish between these two mechanisms; for the mechanism I, $k$ increases linearly with the inhibitor concentration, whereas for the mechanism II the relationship is hyperbolic. We found a linear relationship between $k$ and [I] (**Figure 3**) suggesting that mupirocin binding does not introduce a slow conformational change. The equilibrium rate constants ($k_{on}$ and $k_{off}$) and the inhibitory constant ($K_i$) were obtained by fitting the data to equation 2 and 3, where [I] is the concentration of inhibitor, [S] is the concentration of the substrate, and $K_M$ is corresponding Michaelis value for the substrate.

$$k = k_{\mathrm{off}} + \frac{k_{\mathrm{on}} \times [\mathrm{I}]}{1 + \frac{[\mathrm{S}]}{K_{\mathrm{M}}}} \text{ (eq. 2)}$$

$$K_{\mathrm{i}} = \frac{k_{\mathrm{off}}}{k_{\mathrm{on}}} \text{ (eq. 3)}$$

PmIleRS2 displayed classic fast-on/fast-off competitive inhibition. To determine $K_{\mathrm{i}}$ values with respect to isoleucine and ATP, one substrate was varied from 0.2 – 10 × $K_{\mathrm{M}}$, second substrate was present at 20 × $K_{\mathrm{M}}$ (25 mM ATP or 1 mM isoleucine) and mupirocin was varied from 1 – 20 × $K_{\mathrm{i}}$. Reactions were started by addition of substrate, and enzyme was present at 20 nM. Initial velocity data were fitted directly to global competitive inhibition model using GraphPad Prism.

*Two-step aminoacylation*

Two-step aminoacylation, comprising both the activation of amino acid and its transfer to $tRNA^{Ile}$, was followed at 30 °C in a buffer containing 50 mM Hepes-KOH, pH 7.5, 10 mM $MgCl_2$, 1 mM DTT, 150 mM $NH_4Cl$, 0.008 U/µL TIPP, 0.1 mg/mL BSA, 4 mM ATP, 1 mM Ile, 10-40 µM [$^{32}$P]-$tRNA^{Ile}$, and 1-100 nM enzyme. Steady-state kinetic parameters were determined by varying the corresponding substrate from 0.1 – 10 × $K_{\mathrm{M}}$, while the other substrates were kept at the saturating concentration (5-10 × $K_{\mathrm{M}}$). Reactions were started by adding the substrate (Ile or ATP) and stopped as described for amino acid activation. Quenched reaction mixture was further treated with P1 nuclease [122]. Acceptor activity of $tRNA^{Ile}$ after labeling with [$^{32}$P] was ~78 %. The rate constants were corrected by this factor to account for the proportion of the functional [$^{32}$P]-$tRNA^{Ile}$.

*Single-turnover deacylation (post-transfer editing)*

Post-transfer editing is addressed by measuring enzymatic deacylation of pre-formed misaminoacylated tRNA [125]. We showed previously that the rate of deacylation is faster than the rate of release of the deacylated tRNA from the enzyme [128]. Thus, to determine the deacylation rate constant ($k_{\mathrm{deacyl}}$), one catalytic turnover must be followed. Single-turnover conditions were ensured by using the enzyme in excess over aa-tRNA$^{\mathrm{Ile}}$. Under these conditions the obtained rate constant is not influenced by slow product release. Single-turnover deacylation was performed at 30 °C in a buffer containing 150 mM Hepes-KOH, pH 7.5, 150 mM $NH_4Cl$, 10 mM $MgCl_2$, 1 mM DTT, 10-20 μM enzyme and 200-400 nM renatured aa-[$^{32}$P]-tRNA$^{\mathrm{Ile}}$. Reaction was started by mixing the enzyme with aa-[$^{32}$P]-tRNA$^{\mathrm{Ile}}$. Fast reactions were performed using rapid chemical quench instrument (RQF-3, KinTek Corp). Aa-[$^{32}$P]-tRNA$^{\mathrm{Ile}}$ was prepared as described [122] using the PmIleRS2 post-transfer editing deficient mutant. The data were fitted to single exponential equation $Y = Y_0 + A \times \mathrm{e}^{-k_{\mathrm{deacyl}} \times t}$ where $Y_0$ is the *y* intercept, *A* is amplitude and *t* is time.

*Parallel formation of [$^{32}$P]-AMP and aa-[$^{32}$P]-tRNA$^{Ile}$*

Parallel formation of [$^{32}$P]-AMP and aa-[$^{32}$P]-tRNA$^{\mathrm{Ile}}$ allows determination of the ratio $k_{\mathrm{AMP}}/k_{\mathrm{aa\text{-}tRNA}}$, whose value above 1 diagnoses editing. Reactions were followed at 30 °C in a buffer containing 50 mM Hepes-KOH, pH 7.5, 20 mM $MgCl_2$, 5 mM DTT, 0.004 U/μL, 0.1 mg/mL BSA, 60 μM tRNA$^{\mathrm{Ile}}$ and 200 μM ATP. Reactions were supplemented with either [$^{32}$P]-tRNA$^{\mathrm{Ile}}$ or α-[$^{32}$P]-ATP to monitor synthesis of aa-tRNA$^{\mathrm{Ile}}$ or ATP consumption, respectively. Lower concentration of ATP compared to the aminoacylation reaction was used to increase sensitivity of the assay. The enzymes were present at 100 nM, and concentrations of amino acids were 2 mM Ile, 20 mM Val, 50 mM Nva. The reactions were started by addition of enzyme and quenched as described above (for reactions monitoring aa-[$^{32}$P]-tRNA$^{\mathrm{Ile}}$ formation) or by mixing 1.5 μL reaction mixture with 3 μL 1.5 M formic acid (for reactions monitoring [$^{32}$P]-ATP consumption).

The apparent rate constants for AMP ($k_{AMP}$) and aa-tRNA$^{Ile}$ ($k_{aa\text{-}tRNA}$) formation were determined from the slope of product formation in time divided by the enzyme concentration.

*Protein thermal denaturation*

Thermal denaturation was performed by nanoDSF using Nanotemper Prometheus NT.48 in temperature range from 20 °C do 90 °C with a heating rate of 3 °C/min. Protein was diluted in a buffer containing 20 mM Hepes-KOH, pH 7.5, 50 mM NaCl, 5 mM β-mercaptoethanol and 10 % glycerol to a final concentration of 1 mg/mL and ratio in fluorescence change at 350 nm and 330 nm was used to determine the apparent $T_m$ using PR.Stability Analysis (NanoTemper).

*Total protein isolation and immunoblotting*

Cell pellets were harvested and resuspended in the lysis buffer containing 25 mM Tris-HCl, pH 7.5, 150 mM NaCl, 5 mM $MgCl_2$, 0.1 mM PMSF, 10% (v/v) glycerol, 5 mM β-mercaptoethanol, 2 mg/mL lysozyme, 100 µg/mL DNase I. Cell lysis was assisted by mild sonication. Total protein concentration was estimated using Bradford assay and 30 µg of total proteins were used for immunoblotting. Semi-dry transfer to nitrocellulose membrane was performed using 2117 Multiphor II (LKB) in a buffer containing 25 mM Tris, pH ~ 8.2, 192 mM glycine, 20% (v/v) methanol and 0.1% (w/v) SDS for 1-1.5h at room temperature at constant current 0.8 mA/cm$^2$. The membrane was incubated overnight at 4 °C in 5 % non-fat milk dissolved in TBST (25 mM Tris-HCl, pH 7.5, 150 mM NaCl, 0.1 % (v/v) Tween-20) to block non-specific binding. The membrane was then incubated for 1h at room temperature with anti-His$_6$ (Roche) primary antibodies in the same buffer. Finally, the membrane was incubated in a buffer containing secondary antibodies (anti-Mouse-HRP, Roche) conjugated with HRP for 1 h at room temperature. The membrane was washed three times with TBST after incubation with antibodies. Signals were detected using Amersham ECL Select Western Blotting Detection Reagent (GE Healthcare) followed by exposure to autoradiographic film.

*In gel-digestion of polypeptides*

The same cell extracts used for whole proteome MS analysis were used for SDS-PAGE as well. In-gel digestion of *Coomassie*-stained polypeptides followed standard procedures: The gel slices were destained and dehydrated with acetonitrile. Polypeptides were reduced with DTT (10 mM DTT in 5 mM ammonium bicarbonate, 45 min at 56 °C) and alkylated with iodoacetamide (55 mM iodoacetamide in 5 mM ammonium bicarbonate, 30 min at 25 °C). Gel slices were washed 2 × 20 min with 50 % acetonitrile and 5 % ammonium bicarbonate, dehydrated with acetonitrile, dried under vacuum, and reswollen in trypsin solution (Promega, 12.5 ng/µL trypsin in 20 mM ammonium bicarobonate). The supernatant after overnight digestion was collected and gel slices were extracted with 50 % acetonitrile, 1 % trifluoroacetic acid and 80 % acetonitrile, 1 % trifluoroacetic acid. The extracts and supernatant were pooled, the volume was reduced, and acetonitrile removed by evaporation under vacuum. The samples were desalted on C18 StageTips and prepared for MS analysis as described in the main text. All samples were prepared and analyzed in triplicates.
